# Impaired immune response drives age-dependent severity of COVID-19

**DOI:** 10.1101/2022.04.21.489072

**Authors:** Julius Beer, Stefania Crotta, Angele Breithaupt, Annette Ohnemus, Jan Becker, Benedikt Sachs, Lisa Kern, Miriam Llorian, Nadine Ebert, Fabien Labroussaa, Tran Thi Nhu Thao, Bettina Salome Trueeb, Joerg Jores, Volker Thiel, Martin Beer, Jonas Fuchs, Georg Kochs, Andreas Wack, Martin Schwemmle, Daniel Schnepf

## Abstract

SARS-CoV-2 is a highly contagious respiratory virus and the causative agent for COVID-19. The severity of disease varies from mildly symptomatic to lethal and shows an extraordinary correlation with increasing age, which represents the major risk factor for severe COVID-19^1^. However, the precise pathomechanisms leading to aggravated disease in the elderly are currently unknown. Delayed and insufficient antiviral immune responses early after infection as well as dysregulated and overshooting immunopathological processes late during disease were suggested as possible mechanisms. Here we show that the age-dependent increase of COVID-19 severity is caused by the disruption of a timely and well-coordinated innate and adaptive immune response due to impaired interferon (IFN) responses. To overcome the limitations of mechanistic studies in humans, we generated a mouse model for severe COVID-19 and compared the kinetics of the immune responses in adult and aged mice at different time points after infection. Aggravated disease in aged mice was characterized by a diminished IFN-γ response and excessive virus replication. Accordingly, adult IFN-γ receptor-deficient mice phenocopied the age-related disease severity and supplementation of IFN-γ reversed the increased disease susceptibility of aged mice.

Mimicking impaired type I IFN immunity in adult and aged mice, a second major risk factor for severe COVID-19^2–4^, we found that therapeutic treatment with IFN-λ in adult and a combinatorial treatment with IFN-γ and IFN-λ in aged *Ifnar1^-/-^*mice was highly efficient in protecting against severe disease.

Our findings provide an explanation for the age-dependent disease severity of COVID-19 and clarify the nonredundant antiviral functions of type I, II and III IFNs during SARS-CoV-2 infection in an age-dependent manner. Based on our data, we suggest that highly vulnerable individuals combining both risk factors, advanced age and an impaired type I IFN immunity, may greatly benefit from immunotherapy combining IFN-γ and IFN-λ.

## Introduction

Within two years since its introduction into the human population, SARS-CoV-2 has caused more than 450 million confirmed cases of COVID-19 leading to about 6 million deaths globally as of March 2022 [*WHO coronavirus dashboard*]. Interestingly, the burden of severe disease and mortality is not equally distributed across age groups and shows an extraordinary log-linear correlation with increasing age for individuals older than 30 years^1^. To enable the rational design of effective therapeutics and prevention strategies for vulnerable groups, a better understanding of disease-causing mechanisms is urgently needed.

One common hallmark of severe COVID-19 and advanced age is a diminished and delayed innate immune response affecting the timely production of interferons (IFNs)^5–8^. Type I, II and III IFNs, also called IFN-α/β, IFN-γ and IFN-λ, respectively, are known antiviral cytokines which are rapidly produced by the host upon recognition of viral material. IFNs orchestrate an immediate cell intrinsic innate immune response by upregulating expression levels of interferon-stimulated genes (ISGs) and initiate the subsequent adaptive immune response by the recruitment and activation of immune cells^9–12^. The delayed and diminished IFN response in severe COVID-19 is associated with a late and dysregulated inflammatory gene expression signature^8, 12–14^, likely due to enhanced tissue damage caused by an insufficient control of virus replication. The clinical relevance of a well-functioning IFN response was emphasized by the finding that 3.5 % of patients with life-threatening COVID-19 had genetic defects in genes involved in virus recognition, IFN production and signaling, including *TLR3*, *TBK1*, *IRF3*, *IRF7*, *IFNAR1* and *IFNAR2*^2^. In addition, type I IFN neutralizing antibodies were detected in another 10 % of critically ill COVID-19 patients with a tendency of increased frequency in the elderly^4^. Even though clinical penetrance of neutralizing antibodies against type I IFNs for severe COVID-19 is not complete^15^, they may promote lethal disease progression in up to 20 % of deaths caused by SARS-CoV-2 infection^3^. The markedly lower risk of children to develop severe COVID-19 on the other hand correlates with an increased basal expression level of the pattern-recognition receptors (PRRs) MDA5 and RIG-I leading to a stronger innate antiviral immune response upon SARS-CoV-2 infection compared with adults^16, 17^. In agreement, an early type I IFN response in immune cells was associated with the containment of virus dissemination preventing viral pneumonia^18^. However, despite its important endogenous role, type I IFNs have limited therapeutic potential due to their ability to augment disease late during infection^19^.

In contrast, type III IFNs lack such inflammatory effects^20, 21^ and can be used as potent antiviral treatments, even in the absence of a fully functional type I IFN immunity^22, 23^. A higher IFN-λ to IFN-α/β ratio in critically ill COVID-19 patients correlated with an improved disease outcome, and patients with high expression levels of IFN-λ showed decreased viral loads and an accelerated viral clearance^13^. In line, a small phase II placebo-controlled randomized trial in humans found that treatment of SARS-CoV-2 infected patients with IFN-λ could accelerate viral decline and clearance^24^.

The role of type II IFN during COVID-19 on the other hand is much less clear. Whereas one study reported that the epithelial response to IFN-γ would promote SARS-CoV-2 infection^25^, another one demonstrated significant inhibition of virus replication^26^. NK cells from ambulant COVID-19 patients showed increased production of IFN-γ, whereas NK cells from patients with severe COVID-19 produced only low levels of IFN-γ and TNF^27^. This study further demonstrated that an untimely TGF-β response, a cytokine suppressing IFN-γ mediated functions, was limiting the antiviral activity of NK cells. In agreement, others found that NK cells from severe COVID-19 patients were dysfunctional showing an impairment of antiviral activity that was associated with a diminished production of IFN-γ and TNF^28^.

Despite the vast amount of clinical data available by now, many questions regarding the disease-causing mechanisms in the elderly remain unresolved. Small animal models are essential to overcome the limitations of human sample heterogeneity and availability. However, most clinical isolates of SARS-CoV-2 cannot infect standard inbred mice, with few exceptions only causing asymptomatic infection^29, 30^. Whereas knock-in mice expressing human angiotensin I-converting enzyme 2 (*ACE2*), the receptor for SARS-CoV-2, are permissive for infection with clinical isolates, they do not develop severe or lethal disease^31–33^. Another widely used model that does support severe to lethal disease upon SARS-CoV-2 infection are transgenic mice expressing *ACE2* under the cytokeratin-18 (K18) promotor. While this mouse strain is suitable to test various intervention strategies^34, 35^, its use to study mechanisms of disease is limited, e.g. by artifactual neuroinvasion of the virus due to abundant and non-physiological expression of the viral receptor^36^. To be able to use existing standard inbred mouse strains, including knockout mice, researchers developed mouse-adapted SARS-CoV-2 strains either by *in silico* design followed by reverse genetics^37^, by serial passaging^38–42^ or by a combination of both^43^.

In this study, we describe the generation of a highly pathogenic mouse-adapted SARS-CoV-2 strain (designated SARS-CoV-2 MA20) that dose-dependently causes mild, severe or even lethal disease progression in 8-20-week-old adult C57BL/6 wild type mice. In 36-60-week-old aged C57BL/6 mice, disease severity was strongly enhanced and correlated with (i) the lack of an early and well-coordinated innate and adaptive immune response, (ii) a markedly increased viral load, and (iii) a late inflammatory response. Direct comparison of adult and aged knockout mice showed that defective type I and type II IFN signaling phenocopies enhanced disease progression in aged mice, providing a mechanistic explanation for the age-related increase in disease susceptibility during SARS-CoV-2 infection.

Using adult mice lacking a functional IFN-α/β receptor (*Ifnar1^-/-^*) to mimic an impaired type I IFN immunity^2–4^, we show that prophylactic or therapeutic administration of IFN-λ efficiently protected such lethally infected mice. Nevertheless, IFN-λ treatment alone had limited protective effects in highly vulnerable aged *Ifnar1^-/-^* mice. However, administration of IFN-γ in aged wild type mice reversed the age-dependent enhanced disease phenotype, and a combinatorial treatment with IFN-λ and IFN-γ even protected highly vulnerable aged *Ifnar1^-/-^* mice against lethal disease.

By generating and employing a mouse model for severe COVID-19 we identified an age-dependent impairment of type I and type II IFN responses as a critical pathomechanism that drives the virulence of SARS-CoV-2 in aged hosts. This novel insight was successfully translated into an immunomodulatory treatment strategy that prevented SARS-CoV-2-induced lethality in a highly susceptible disease model that mimics impaired type I IFN immunity and advanced age.

## Results

### Type I and type III IFNs synergize to limit SARS-CoV-2 replication and protect aged mice against symptomatic disease

To dissect the individual and combinatorial role of type I and type III IFNs in limiting SARS-CoV-2 replication, we compared the replication kinetics of a mouse-adapted but largely non-virulent SARS-CoV-2 strain (SARS-CoV-2 MA^37^) in IFN receptor-deficient and C57BL/6 wild type (WT) mice. Three days post infection (d p. i.) we found about 10-fold increased viral loads in lungs and upper airways of mice lacking functional type I (*Ifnar1^-/-^*) or type III IFN receptors (*Ifnlr1^-/-^*) (**Figure 1A** and **Figure S1A**). By day 5 p. i. most WT mice had cleared the virus, whereas *Ifnar1^-/-^* and especially *Ifnlr1^-/-^* mice continued to have high viral titers in their lungs (**Figure 1A**). Conversely, prophylactic or therapeutic administration of either IFN-α_B/D_^22^ to *Ifnlr1^-/-^* or IFN-λ1/3^44^ to *Ifnar1^-/-^* mice reduced lung viral loads on day 3 p. i. by several orders of magnitude (**Figure 1B** and **Figure S1B**). Combinatorial loss of type I and type III IFN signaling in *Ifnar1^-/-^Ifnlr1^-/-^* mice led to excessive replication and prolonged persistence of the virus in upper airways and lungs (**Figure 1A** and **Figure S1A**). Immunohistochemical analyses confirmed increased virus replication and prolonged virus persistence in *Ifnar1^-/-^Ifnlr1^-/-^* compared with WT mice (**Figure S1C-D**), mainly affecting the bronchial epithelium and only scarcely alveolar epithelial cells. Lung tissue damage (**Figure S1E**) and necrotizing bronchitis (**Figure S1F**) followed the pattern of increased and prolonged virus replication in *Ifnar1^-/-^ Ifnlr1^-/-^*mice. However, despite increased and prolonged viral replication and tissue damage, we did not observe weight loss or other signs of disease in adult mice lacking type I and/or type III IFN responses (**Figure 1C**). In contrast, using aged animals in an identical infection setting, we observed significantly increased weight loss in *Ifnar1^-/-^*, *Ifnlr1^-/-^* and *Ifnar1^-/-^Ifnlr1^-/-^* mice compared with age-matched WT controls (**Figure 1D**), sporadically even leading to lethal disease progression in case of *Ifnar1^-/-^Ifnlr1^-/-^* mice (**Figure S1G**). Of note, we did not observe increased weight loss in aged WT mice compared with adult controls (**Figure 1C and D**), indicating that a combination of advanced age and impaired type I/III IFN immunity is required to result in symptomatic disease during infection with the SARS-CoV-2 MA strain.

**Figure 1.**
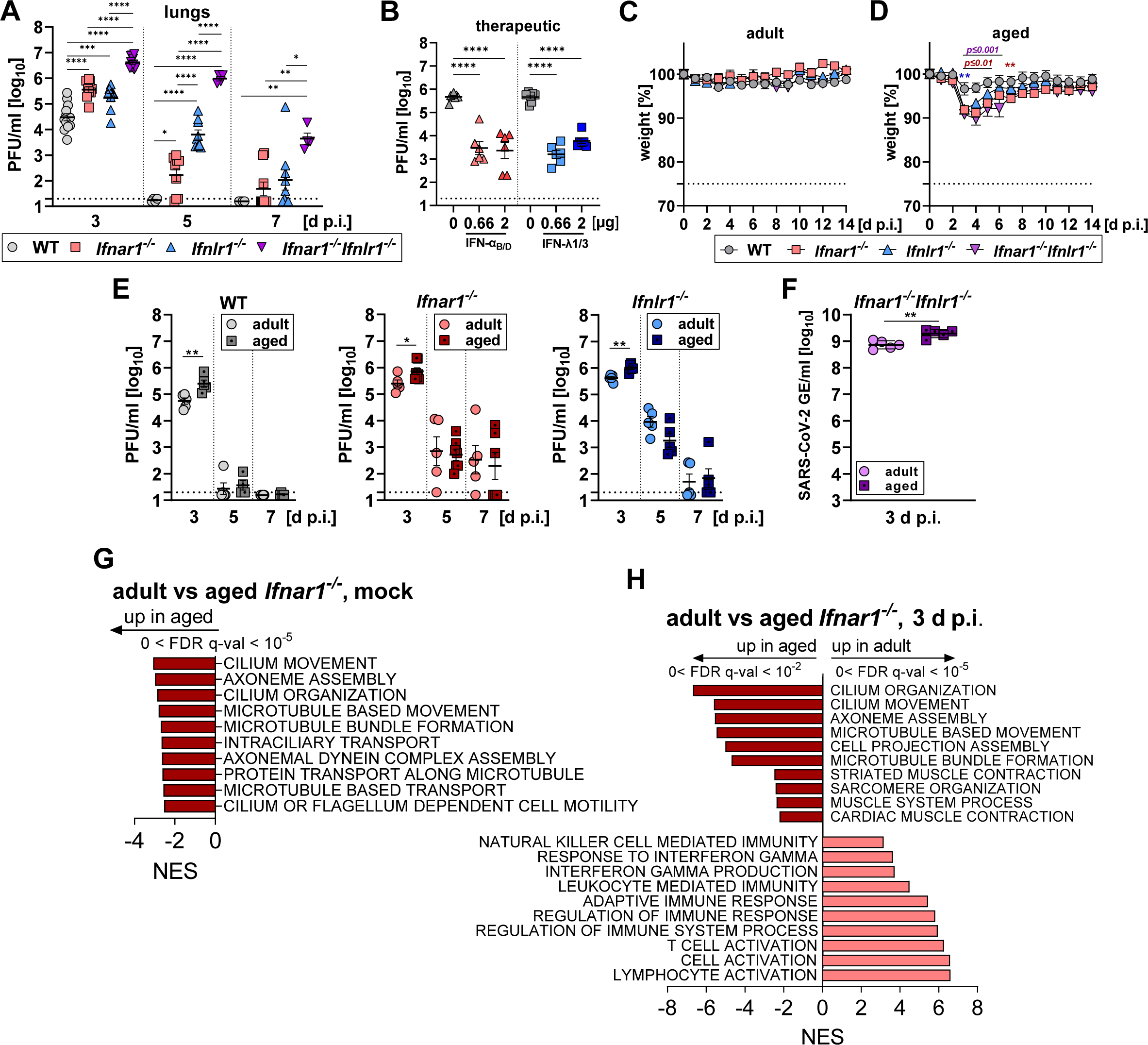
Increased and prolonged replication of SARS-CoV-2 MA causes symptomatic disease in aged mice lacking type I and/or type III IFN receptors which correlates with an impaired immune response. **A)** Groups of adult (8-18-week-old) mice of the indicated genotypes were infected with 10^5^ PFU SARS-CoV-2 MA. Lungs were harvested at the indicated time points and viral loads determined by plaque assay on Vero E6 cells. Data pooled from five independent experiments. Symbols represent individual mice (n=4-13 per group) and bars indicate mean ± SEM. Dashed line indicates detection limit. \**P*≤0.05, \*\**P*≤0.01, \*\*\**P*≤0.001, \*\*\*\**P*≤0.0001, one-way ANOVA with Tukey’s multiple comparisons test. **B)** Groups of 16-24-week-old *Ifnlr1*^-/-^ (triangles) and *Ifnar1*^-/-^ mice (squares) were treated intranasally with the indicated doses of IFN-α_B/D_ or IFN-λ1/3, respectively, or mock-treated, one day after infection with 10^5^ PFU SARS-CoV-2 MA. Lung viral loads on day 3 p. i. were determined by plaque assay on Vero E6 cells. Data from a single experiment are shown. Symbols represent individual mice (n=6-7 per group) and bars indicate mean ± SEM. Dashed line indicates detection limit. \*\*\*\**P*≤0.0001, one-way ANOVA with Tukey’s multiple comparisons test. **C-D)** Groups of adult (10-15-week-old; n=7-8) (**C**) or aged (36-60-week-old mice; n=10-19) (**D**) of the indicated genotypes were infected with 10^5^ PFU SARS-CoV-2 MA. Signs of disease and weight loss were monitored for 14 days. Data from a single experiment are shown in (**C**) and pooled data from two independent experiments are shown in (**D**). Symbols represent mean ± SEM. Dashed line indicates experimental endpoint due to animal welfare. Two-way ANOVA with Tukey’s multiple comparisons test comparing WT with *Ifnar1^-/-^* (red), *Ifnlr1^-/-^* (blue) and *Ifnar1^-/-^Ifnlr1^-/-^* (purple), \*\**P*≤0.01. **E)** Groups of adult (8-12-week-old) or aged (40-60-week-old) mice (n=4-7) of the indicated genotypes were infected with 10^5^ PFU SARS-CoV-2 MA. Lungs were harvested at the indicated time points and viral loads determined by plaque assay on Vero E6 cells. Data pooled from four independent experiments are shown. Symbols represent individual mice and bars indicate mean ± SEM. Dashed line indicates detection limit. \**P*≤0.05, \*\**P*≤0.01, unpaired t test. **F)** Groups of adult (10-week-old) or aged (48-54-week-old) *Ifnar1^-/-^Ifnlr1^-/-^*mice (n=5) of the indicated genotypes were infected with 10^5^ PFU SARS-CoV-2 MA. Lungs were harvested on day 3 p. i. and viral RNA levels (*E* gene) were quantified by RT-qPCR. Data from a single experiment are shown. Symbols represent individual mice and bars indicate mean ± SEM. \*\**P*≤0.01, unpaired t test. **G-H**) Groups of adult (12-week-old) or aged (44-week-old) *Ifnar1^-/-^*mice (n=5) were mock-infected (**G**) or infected with 10^5^ PFU SARS-CoV-2 MA (**H**). Lungs were harvested on day 3 p. i. and processed for RNA sequencing. Gene Set Enrichment Analysis (GSEA) of ranked genes indicating strong enrichment for immune cell activation pathways in infected adult compared with aged mice. Negative and positive normalized enrichment scores (NES) indicate enrichment in aged and adult *Ifnar1^-/-^* mice, respectively. False Discovery Rate (FDR) < 10^-2^ for all pathways shown.

Taken together these data demonstrated that type I and type III IFNs synergize to limit excessive SARS-CoV-2 replication, to expedite virus clearance and to protect against symptomatic disease in aged mice.

### Advanced age correlates with increased viral loads and diminished immune responses

To gain more insights into the age-dependent disease phenotype of SARS-CoV-2 MA, we compared tissue sections from upper airway and lung samples of infected adult and aged *Ifnar1^-/-^* mice. Three days p. i., we detected increased antigen load in upper airways and lungs as well as enhanced bronchial necrosis in aged *Ifnar1^-/-^* mice (**Figure S1H-J**). Comparing viral growth kinetics in adult and aged WT, *Ifnar1^-/-^*, *Ifnlr1^-/-^*, and *Ifnar1^-/-^Ifnlr1^-/-^* mice, we confirmed the age-dependent increase in virus replication at 3 d p. i., irrespectively of genotype (**Figure 1E-F** and **Figure S1K**). To identify impaired antiviral or enhanced pro-viral pathways facilitating virus replication in aged mice that are independent of type I/III IFN signaling, we performed transcriptome analyses using lung samples of infected or mock-treated adult and aged *Ifnar1^-/-^*mice. Gene Set Enrichment Analyses (GSEA) comparing mock-treated (**Figure 1G**) or infected (**Figure 1H**) adult and aged *Ifnar1^-/-^* mice both identified an age-related increase in pathways involved in the function of ciliated cells, possibly suggesting age-dependent differences in the cellular composition of the lung. Whereas no pathways were significantly enriched in uninfected adults compared to aged mice, lung tissue samples from infected adult mice showed a significant enrichment in pathways involved in the production and response to IFN-γ, NK cell-mediated immunity, immune cell activation and adaptive immune responses (**Figure 1H**) which indicated a versatile and robust immune response in adult animals. The disruption of a timely and well-coordinated innate and adaptive immune response in aged mice upon SARS-CoV-2 infection could well explain impaired virus control, ultimately leading to enhanced disease progression.

These data demonstrate that the age-dependent increase in virus replication is independent of type I and type III IFN signaling, but correlates with an age-related impaired immune response affecting IFN-γ production, NK cell mediated immunity and general immune cell activation.

### Rapid host adaptation by serial passaging in type I/III IFN receptor-deficient C57BL/6 mice

To study the age-related pathophysiology of SARS-CoV-2-induced disease in more detail and to test possible intervention strategies, we generated a mouse model resembling severe COVID-19 by serially passaging the SARS-CoV-2 MA strain *in vivo*. In total, we performed four independent passaging series, two in C57BL/6 WT (WT A and B) and two in *Ifnar1^-/-^Ifnlr1^-/-^* mice (DKO A and B) (**Figure 2A**). From passage 10 onwards, *Ifnar1^-/-^Ifnlr1^-/-^* but not WT mice were losing increasing amounts of their initial body weight (**Figure 2B**). Whereas viral titers in lungs remained relatively stable until passage 20, viral loads in the upper airways increased from passage 14 onwards for series WT B, DKO A and DKO B (**Figure S2A-B**). To identify which passaging series contained pathogenic variants, we infected groups of C57BL/6 WT mice with passage 20 (P20) lung homogenates containing 10^4^ plaque forming units (PFU) of virus and followed the course of disease and survival rates (**Figure 2C**). Virus variants derived from passaging series WT A and B did not induce severe signs of disease, whereas P20 homogenates from passaging series DKO A and DKO B induced severe weight loss and even 40 % lethality in adult C57BL/6 WT mice in case of DKO A P20. Using plaque-purified (PP) virus stocks derived from DKO A and DKO B P20 lung homogenates, we confirmed the successful generation of a pathogenic mouse-adapted SARS-CoV-2 variant that emerged in passaging series DKO A (**Figure S2C**). Virus genome sequencing revealed that this variant, named SARS-CoV-2 MA20 (MA20), acquired eight additional amino acid changes compared with the parental SARS-CoV-2 MA strain (**Figure 2D**), three in S (T250A, K417N and Q493H), one in M (T7I) and four in ORF1ab (A1997V, T3058I, D4165Y and T4174A) that translate into A1179V in nsp3, T295I in nsp4, and D25Y and T34A in nsp9. Of note, identical or similar amino acid substitutions such as the Q493H and K417N in the spike protein are also present in other mouse-adapted SARS-CoV-2 variants^39, 40, 43^ or circulating variants of concern (VOCs) including B.1.1.529 (Omicron) (**Figure S2D-E**). Despite its increased pathogenicity in mice, SARS-CoV-2 MA20 replication was strongly attenuated in human Calu-3 and simian Vero E6 cells (**Figure 2E** and **Figure S2F**). In addition, MA20 is even better neutralized by sera from vaccinated humans compared with the B.1.617.2 (Delta) variant (**Figure S2G**). Using doses from 10^2^-10^4^ PFU MA20, we could model mild, moderate, or even lethal SARS-CoV-2-induced disease progression in adult C57BL/6 WT mice (**Figure 2F**). By infecting age- and sex-matched C57BL/6, BALB/c and 129/sv mice with 10^3^ PFU MA20, we found that BALB/c mice were highly susceptible with a survival rate of only 30 %, that 129/sv mice were mostly resistant to disease, and that C57BL/6 showed intermediate susceptibility with substantial weight loss but a survival rate of 90 % (**Figure 2G**).

**Figure 2.**
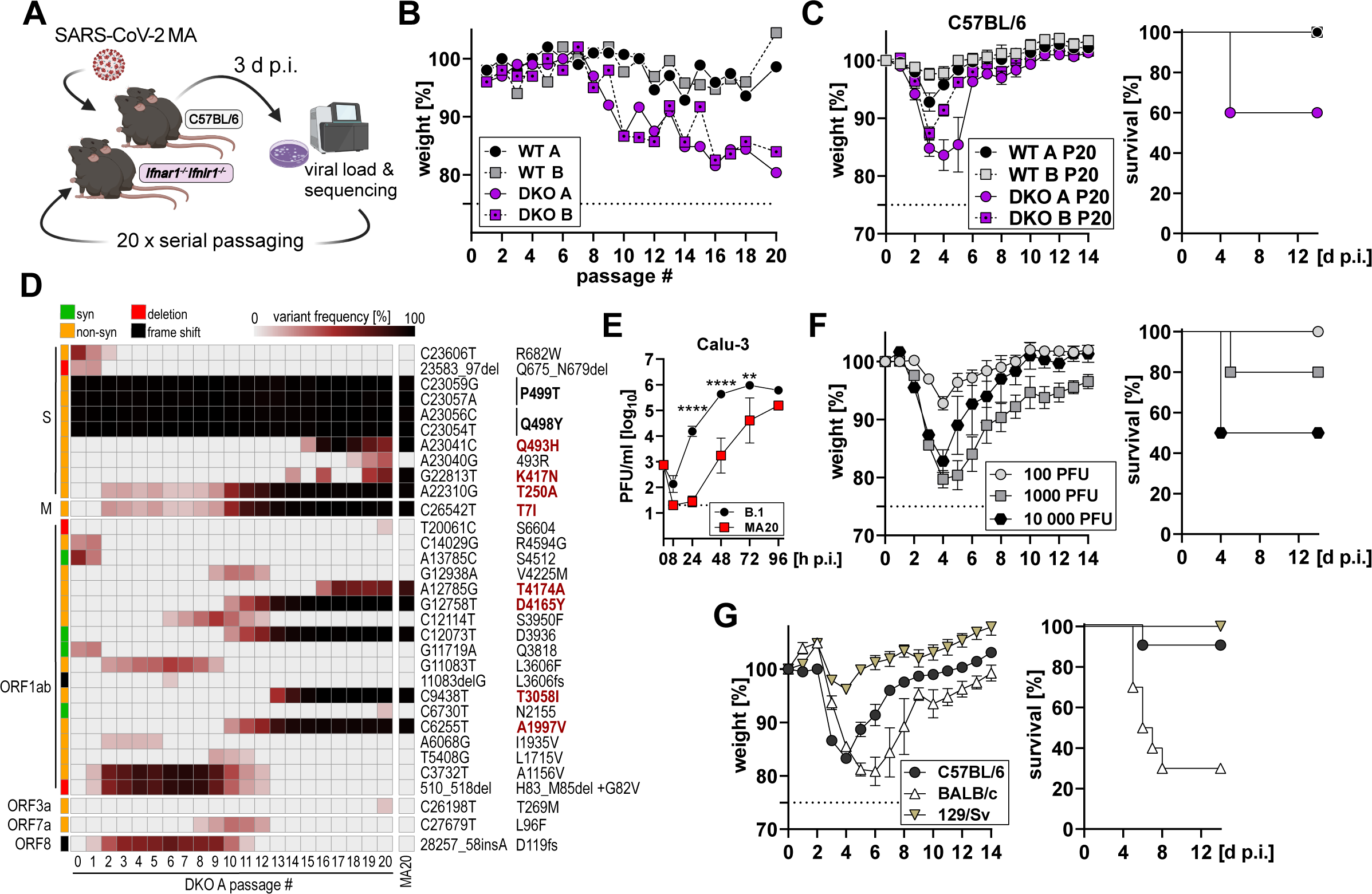
Serial passaging of SARS-CoV-2 MA in *Ifnar1^-/-^Ifnlr1^-/-^* C57BL/6 mice yielded a highly virulent virus variant. **A)** Schematic overview of serial passaging of SARS-CoV-2 MA in C57BL/6 (WT A and B) and B6-*Ifnar1^-/-^Ifnlr1^-/-^* mice (DKO A and B). Mice were initially infected with 10^5^ PFU SARS-CoV-2 MA. Serial passaging was performed by infecting mice with lung homogenates containing 10^3^-10^5^ PFU. Lungs were harvested on day 3 post infection. **B)** Weight loss of C57BL/6 (WT A and B) and B6-*Ifnar1^-/-^Ifnlr1^-/-^*mice (DKO A and B) per passage on day 3 post infection. Symbols represent individual mice. Dashed line indicates experimental endpoint due to animal welfare. **C)** Weight loss (left panel) and survival (right panel) of adult C57BL/6 WT mice (8-10-week-old; n=5 per group) infected with diluted passage 20 (P20) lung homogenates derived from passaging series WT A, WT B, DKO A and DKO B containing 10^4^ PFU. Data from a single experiment are shown. Symbols represent mean ± SEM. Dashed line indicates experimental endpoint due to animal welfare. **D)** Variant frequency plot from next-generation sequencing results for passaging series DKO A and the plaque-purified SARS-CoV-2 MA20 virus stock. Variant frequencies are shown in comparison to Wuhan-Hu-1 (NC_045512.2). Amino acid changes present in SARS-CoV-2 MA20 are indicated in bold, changes in comparison to SARS-CoV-2 MA are highlighted in red. **E)** Comparative growth curves of SARS-CoV-2 strains B.1 and MA20 on Calu-3 cells infected with an MOI of 0.001. Virus replication was quantified by plaque assay on Vero E6 cells. Data from a single experiment performed in duplicates are shown. Dashed line indicates detection limit. \*\**P*≤0.01, \*\*\*\**P*≤0.0001, two-way ANOVA with Tukey’s multiple comparisons test. **F)** Weight loss (left panel) and survival (right panel) of adult C57BL/6 mice (10-14-week-old; n=5-10 per group) infected with the indicated dose of SARS-CoV-2 MA20. Pooled data from three independent experiments are shown. Symbols represent mean ± SEM. Dashed line indicates experimental endpoint due to animal welfare. **G)** Weight loss (left panel) and survival (right panel) of adult C57BL/6, 129/Sv or BALB/c mice (10-week-old; n=10 per group) infected with 10^3^ PFU SARS-CoV-2 MA20. Data from a single experiment are shown. Symbols represent mean ± SEM. Dashed line indicates experimental endpoint due to animal welfare.

Next, we performed virus growth kinetics in groups of C57BL/6 WT mice using 10^3^ PFU of MA20 and monitored virus shedding via the nostrils (**Figure 3A**) and virus replication in upper airways (**Figure 3B**) and lungs (**Figure 3C**). Infection-induced gene expression levels of inflammatory cytokines such as *Il6* and *Tnf*, type I (*Ifna4* and *Ifnb*) and type III IFNs (*Ifnl2/3*) as well as IFN-stimulated genes (ISGs) such as *Mx1*, *Isg15* and *Stat1* peaked simultaneously with peak viral loads on day two p. i. in lungs and upper airways (**Figure 3D** **and Figure S2H**). By day seven p. i., no infectious virus could be detected anymore which was in line with histopathological findings that viral antigens were mostly cleared by day 7 (**Figure 3E-F**). Despite rapidly decreasing lung viral loads (**Figure 3C**), lung tissue damage remained at high scores until day 7 (**Figure 3G**). Compared with the less virulent SARS-CoV-2 MA strain, the highly pathogenic SARS-CoV-2 MA20 variant caused a more widespread infection of the lung tissue as indicated by increased antigen-positive areas (**Figure S1I and 3F**; about 10 % mean antigen detection in SARS-CoV-2 MA infected adult *Ifnar1^-/-^* 3 d p. i., compared with about 67% mean antigen detection in SARS-CoV-2 MA20 infected adult WT mice 3 d p. i.). Besides the bronchial epithelium, mainly alveolar epithelial cells were found to be virus-positive, in particular type 2 pneumocytes. In line with an acute viral pneumonia, lung lesions were characterized by necrotizing bronchitis, most severely affecting mice on day 3 p. i. (**Figure 3H**). The extent and severity decreased over time, but bronchial lesions were still detectable until day 7 p. i. in all animals analyzed. Perivascular infiltrates comprised mainly of neutrophils and few lymphocytes. Focal to multifocal necrosis of the alveolar epithelium was associated with minimal to moderate alveolar infiltrates. Tissue regeneration in some animals could be detected as early as 3 d p. i. and consistently increased until day 7 p. i. (**Figure 3I**), as indicated by bronchial epithelial hypertrophy and hyperplasia as well as type 2 pneumocyte hyperplasia. In single cases, we found atypical multinucleated (syncytial) cells, increased mucus production and endotheliitis. Numerous examples for leukocyte rolling were found in blood vessels of infected animals, indicating endothelial and/or immune cell activation. Interstitial infiltrates were rarely detected. Neither vasculitis nor diffuse alveolar damage was detected. Examining other organs by RT-qPCR for the presence of viral RNA, we found low levels in brain and spleen samples and infrequently positive heart, liver, intestine and kidney samples, indicating that replication of SARS-CoV-2 MA20 is mainly restricted to the upper and lower respiratory tract in adult mice (**Figure 3J**).

**Figure 3.**
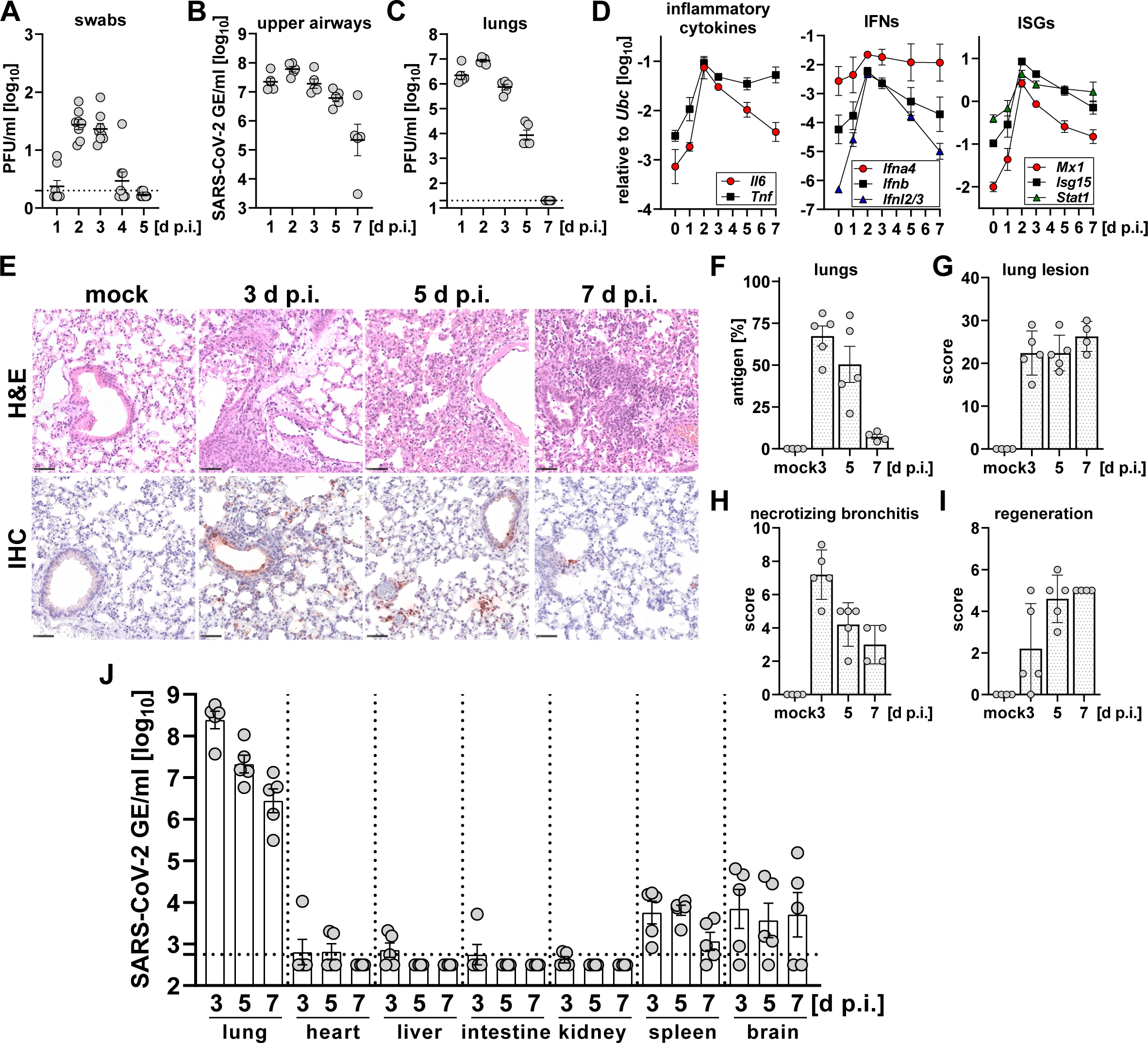
SARS-CoV-2 MA20 efficiently replicates in upper airways and lungs of adult C57BL/6 mice inducing innate immune responses and tissue damage. **A-J**) Adult C57BL/6 mice (8-17-week-old; n=4-8 per group) were mock-treated or infected with 10^3^ PFU SARS-CoV-2 MA20. Samples were collected at the indicated time points. (**A**) Nasal swabs were taken at the indicated time points and viral loads determined by plaque assay on Vero E6 cells. Symbols represent individual mice and bars indicate mean ± SEM. Dashed line indicates detection limit. (**B**) Viral replication in upper airways was quantified as SARS-CoV-2 genome equivalents per ml by measuring expression levels of the viral gene *E* by RT-qPCR at the indicated time points. Symbols represent individual mice and bars indicate mean ± SEM. (**C**) Viral load in lungs was determined by plaque assay on Vero E6 cells at the indicated time points. Symbols represent individual mice and bars indicate mean ± SEM. Dashed line indicates detection limit. (**D**) Gene expression levels of *Il6*, *Tnf*, *Ifna4*, *Ifnb*, *Ifnl2/3*, *Mx1*, *Isg15* and *Stat1* in lungs were determined relative to *Ubc* by RT-qPCR. Symbols represent mean ± SD. **E-I**) Mice were prepared for histological analyses by cardiac perfusion at the indicated time points. (**E**) Representative pictures for histopathology and immunohistochemistry are shown. Bar indicates 50 µm. Antigen (**F**) and histopathologic lesion scores for lungs (**G**) and necrotizing bronchitis (**H**) and regeneration scores (**I**) were quantified as described in materials and methods section. (**J**) Viral RNA levels in lung, heart, liver, intestine, kidney, spleen and brain was quantified as SARS-CoV-2 genome equivalents per ml by measuring expression levels of the viral gene *E* by RT-qPCR at the indicated time points. Symbols represent individual mice and bars indicate mean ± SEM. Dashed line indicates detection limit.

These data show that serial passaging in *Ifnar1^-/-^Ifnlr1^-/-^* mice facilitated rapid host adaption, which resulted in the highly virulent SARS-CoV-2 MA20 variant that can be used to model mild, severe or even lethal COVID-19 in standard inbred mice.

### Enhanced disease progression in aged mice correlates with a diminished immune response leading to insufficient control of virus replication

Disease severity and risk of death due to COVID-19 shows a log-linear correlation with advanced age in humans^1^. Correspondingly, aged mice showed a massively enhanced disease phenotype and increased lethality upon infection with MA20 compared with adult counterparts (**Figure S3A**). To mechanistically address the age-dependent enhanced disease progression, we chose infection conditions which cause a comparable weight loss from which adult but not aged mice could recover (**Figure 4A**) and measured virus replication kinetics, determined systemic dissemination of viral material, assessed lung tissue damage and compared kinetics of the age-dependent immune response profiles in infected lungs. Between day 3 and 5 p. i., virus replication in upper airways (**Figure 4B**) and lungs (**Figure 4C**) of aged mice was found to be increased by one to two orders of magnitude compared with genetically identical adult controls, demonstrating an age-dependent impairment of virus control. Immunohistochemistry of infected lung sections confirmed significantly increased antigen scores in aged mice compared with adult counterparts (**Figure 4D-E**). For both, adult and aged mice, viral antigen was found abundantly in the bronchial and alveolar epithelium, in particular in type 2 pneumocytes. Intriguingly, histopathological analyses revealed that overall lung lesion scores were comparable between both age groups (**Figure 4F**), even though signs of disease were much more pronounced in MA20-infected aged mice. Acute necrotizing bronchitis showed only a slight tendency to be increased in aged mice (**Figure 4G**), and tissue regeneration indicated by bronchial epithelial hypertrophy and hyperplasia as well as type 2 pneumocyte hyperplasia appeared to be less prominent in aged animals (**Figure 4H**). Investigating the potential systemic dissemination of virus material, we found viral RNA levels to be significantly increased in heart, liver, kidney and brain samples of aged mice compared with adult controls on day 4 and 5 p. i. (**Figure 4I**). However, using immunohistochemistry and RNA in situ hybridization methods, no viral antigen or RNA could be detected in heart and brain samples (data not shown). Furthermore, histology of the brain and heart revealed no abnormalities and neither immunohistochemistry for T-cells (CD3) nor microglia/macrophages (Iba-1) identified inflammatory infiltrates or microglial reaction (data not shown).

**Figure 4.**
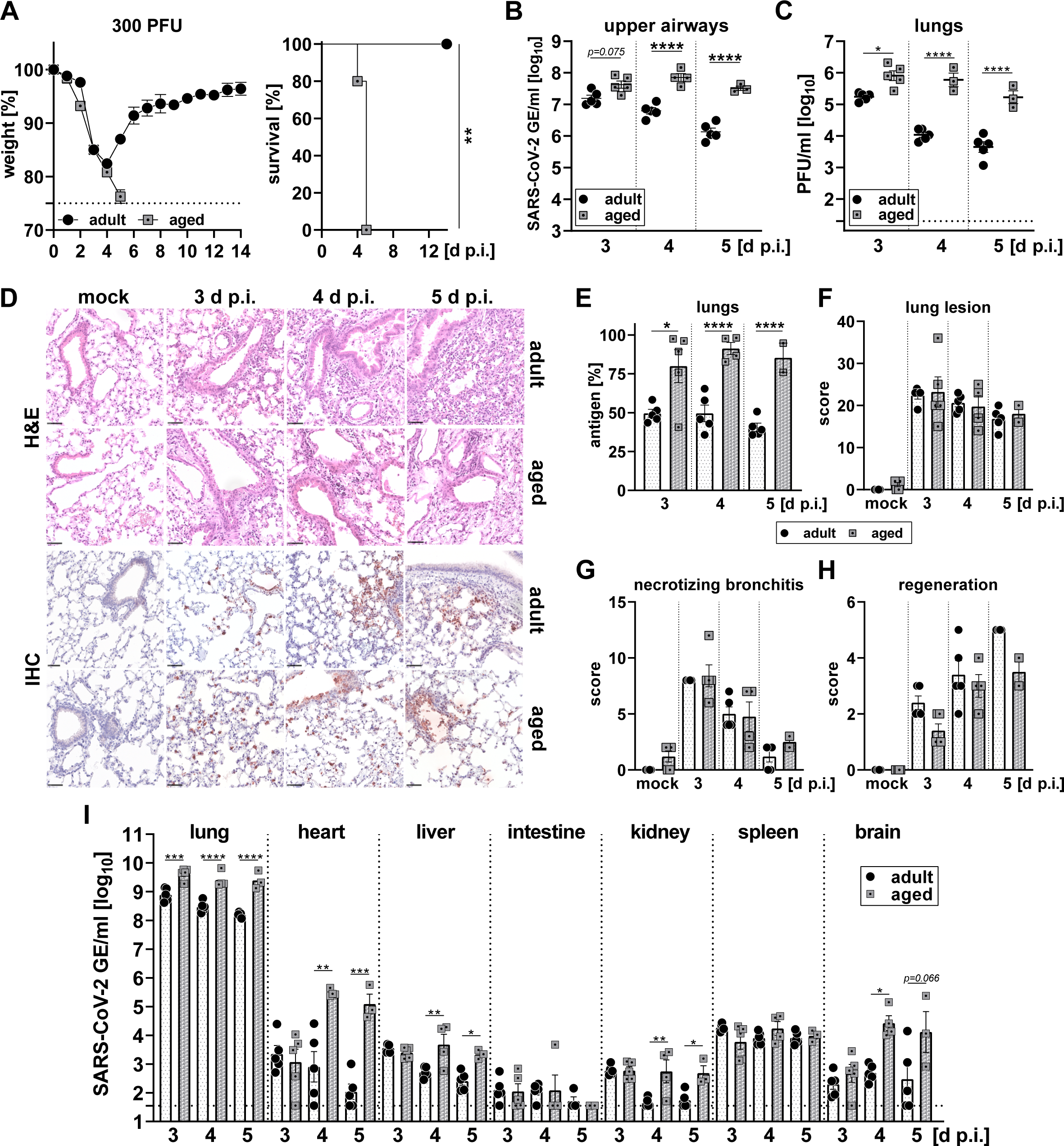
Enhanced susceptibility of aged mice to SARS-CoV-2-induced disease correlates with increased viral load. **A-I**) Groups of adult or aged (10-week- or 40-week-old) C57BL/6 mice were infected with 300 PFU SARS-CoV-2 MA20. Subgroups of mice were sacrificed at the indicated time points to quantify virus replication and to evaluate virus-induced tissue damage. **A)** Weight loss (left panel) and survival (right panel) was monitored for 14 days post infection. Data from a single experiment are shown. Symbols represent mean ± SEM. Dashed line indicates experimental endpoint due to animal welfare. Survival: \*\**P*≤0.01, Log-rank (Mantel-Cox) test, n = 5 per group. **B)** Viral replication in upper airways was quantified as SARS-CoV-2 genome equivalents per ml by measuring expression levels of the viral gene *E* by RT-qPCR at the indicated time points. Data from a single experiment are shown. Symbols represent individual mice and bars indicate mean ± SEM. \**P*≤0.05, \*\*\*\**P*≤0.0001, one-way ANOVA with Tukey’s multiple comparisons test, n = 3-5 per group. **C)** Viral load in lungs was determined by plaque assay on Vero E6 cells at the indicated time points. Data from a single experiment are shown. Symbols represent individual mice and bars indicate mean ± SEM. Dashed line indicates detection limit. \**P*≤0.05, \*\*\*\**P*≤0.0001, one-way ANOVA with Tukey’s multiple comparisons test, n = 3-5 per group. **D-H**) Groups of mice (n = 2-5) were prepared for histological analyses by cardiac perfusion at the indicated time points. (**D**) Representative pictures for histopathology and immunohistochemistry are shown. Bar indicates 50 µm. Antigen (**E**) and histopathologic lesion scores for lungs (**F**), necrotizing bronchitis (**G**) and regeneration scores (**H**) were quantified as described in materials and methods section. **I**) Viral RNA in lungs, heart, liver, intestine, kidney, spleen and brain was quantified as SARS-CoV-2 genome equivalents per ml by measuring expression levels of the viral gene *E* by RT-qPCR at the indicated time points. Symbols represent individual mice and bars indicate mean ± SEM. Dashed line indicates detection limit. \**P*≤0.05, \*\**P*≤0.01, \*\*\**P*≤0.001, one-way ANOVA with Tukey’s multiple comparisons test, n = 3-5 per group.

Next, we compared the kinetics of transcriptional responses in MA20-infected lungs of adult and aged mice. Principal component analysis of the lung transcriptome showed that uninfected adult and aged mice closely clustered together, indicating that basal gene expression profiles in uninfected lungs of adult and aged mice were rather similar (**Figure 5A**). In contrast, transcriptional profiles of infected lung samples clearly diverged dependent on the respective age group for all time points p. i. analyzed, demonstrating a drastically different transcriptional response to SARS-CoV-2 infection between the two age groups (**Figure 5A**). Gene Set Enrichment Analyses (GSEA) comparing each time point p. i. between adult and aged mice revealed that adult animals mounted a rapid and versatile innate and adaptive immune response. In contrast, the immune response of aged animals was delayed, reduced in pathways leading to adaptive immunity, and mainly pro-inflammatory. From 3 d p. i. onwards, innate immune pathways involving IFN-γ signaling and NK cell activity but also adaptive cellular and humoral immune responses were significantly enriched in adult animals compared with aged controls (**Figure 5B**). By contrast, in aged animals primarily pro-inflammatory pathways driven by IL-6, IL-1 and type I IFN were found to be significantly enriched upon day 4 p. i. compared with samples derived from adult animals. Pairwise comparison of infected to mock lung samples from adult or aged animals at different time points post infection using the Ingenuity Pathway Analysis (IPA) confirmed that adult animals were mounting a rapid and well-orchestrated innate and adaptive immune response, characterized by the initiation of PRR signaling, NK cell activation, and the production and response to Th1/Th2 cytokines (**Figure 5C**). In contrast, aged animals showed a reduced, delayed and more pro-inflammatory response. In addition to IL-6- and IL-1-driven pathways, IPA also identified active processes of pulmonary fibrosis and hypoxia-induced gene regulation in infected lung tissue of aged mice (**Figure 5C**). Intriguingly, aged animals also showed an early and strong IL-10 response^45, 46^ which further emphasizes the imbalanced early immune response in aged animals and might explain the lack of a potent immune response initiated by IFN-γ and other immune activating cytokines as observed in adult mice. Taken together these data demonstrate that the age-dependent increase in disease susceptibility upon SARS-CoV-2 infection correlates with an impaired virus control due to imbalanced and insufficient innate and adaptive immune responses.

**Figure 5.**
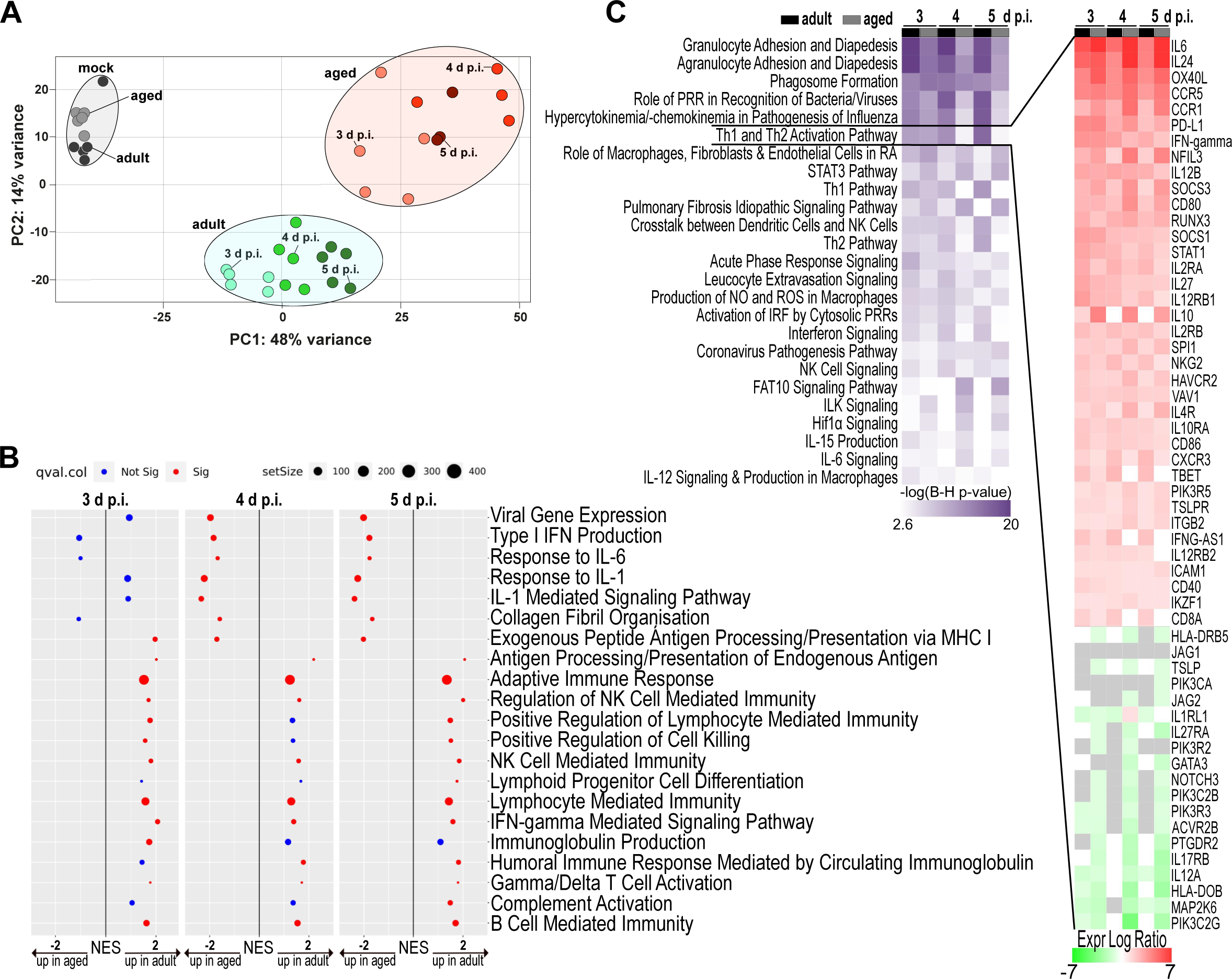
Immune response in SARS-CoV-2 infected aged mice is delayed, diminished and dysregulated. **A-C**) Groups of adult or aged (10-week- or 40-week-old) C57BL/6 mice (n=3-5 per group) were infected with 300 PFU SARS-CoV-2 MA20. Lungs were harvested at the indicated time points, processed and subjected to RNA sequencing. **A)** Principal component analysis (PCA) plot of RNA-sequencing data obtained from total lungs of the indicated groups of mice. **B)** Gene Set Enrichment Analyses (GSEA) comparing adult and aged mice at each time point post infection. Gene set pre-ranked analyses were carried out using the C5 gene ontology (GO) gene set collection in the Molecular Signatures Database (MSigDB). **C)** Heat map for significant differences in Canonical Pathways as defined by Ingenuity Pathway Analysis (IPA). Six pairwise comparisons between infected lung samples and their respective mock controls, from adult or aged animals, at the indicated days p. i., are shown (fold change >1.5, *p*adj < 0.05). The Benjamini-Hochberg method of multiple testing correction was used to calculate -log(p-values). Up- and down-regulated transcripts across the different pairwise comparisons in the “Th1 and Th2 pathway“ are shown as heat map.

### Combinatorial defects in type I and type II IFN signaling phenocopies age-dependent disease susceptibility

Using 300 PFU of the highly virulent MA20 strain we determined the disease susceptibility of adult mice lacking functional type I and/or type III IFN systems. Single knockout mice deficient in type I or type III IFN-mediated responses had comparable survival rates as WT mice but suffered from increased and prolonged weight loss. In contrast, mice lacking both systems rapidly lost weight, and most animals reached experimental endpoints and had to be euthanized (**Figure 6A**). As the disease course of adult *Ifnar1^-/-^Ifnlr1^-/-^* mice closely resembled the one in aged WT mice (**Figure 4A**), we performed low-dose infection experiments with MA20 in adult and aged mice of matching genotypes to assess whether the age-dependent increase in disease susceptibility was caused by an impaired IFN response. Using 30 PFU of MA20, we detected a significantly enhanced disease progression in aged WT, and in addition significantly increased lethality in aged *Ifnar1^-/-^*, *Ifnlr1^-/-^* and *Ifnar1^-/-^Ifnlr1^-/-^* mice compared with their respective adult controls (**Figure 6B-E**). These results indicated that individual or combinatorial loss of type I and type III IFN signaling does not recapitulate the mechanisms underlying increased disease severity in aged mice. Because GSEA and IPA of SARS-CoV-2 infected lung samples both identified an age-dependent reduction in the IFN-γ-mediated immune response to infection, we also assessed the contribution of an impaired type II IFN response to the age-related phenotype. Interestingly, a marked weight loss within 4 d p. i. was only observed in aged but not adult WT, *Ifnar1^-/-^*, *Ifnlr1^-/-^*and *Ifnar1^-/-^Ifnlr1^-/-^* mice (**Figure 6B-E**), whereas weight loss of adult *Ifngr1^-/-^* mice within the first 4 days of infection was comparable to that of aged *Ifngr1^-/-^* mice (**Figure 6F**). This supports our previous findings obtained by transcriptional profiling (**Figure 1H and 5B-C**) that suggested a relevant contribution of an early and potent IFN-γ response to prevent SARS-CoV-2-induced disease in adult mice. Further supporting this hypothesis, initial weight loss and survival rates of infected adult and aged *Ifnar1^-/-^Ifngr1^-/-^* were found to be nearly identical (**Figure 6G**). In addition, adult *Stat1-*deficient mice lacking the ability to respond to any type of IFN, were equally susceptible to SARS-CoV-2-induced disease and lethality as their aged counterparts (**Figure 6H**).

**Figure 6.**
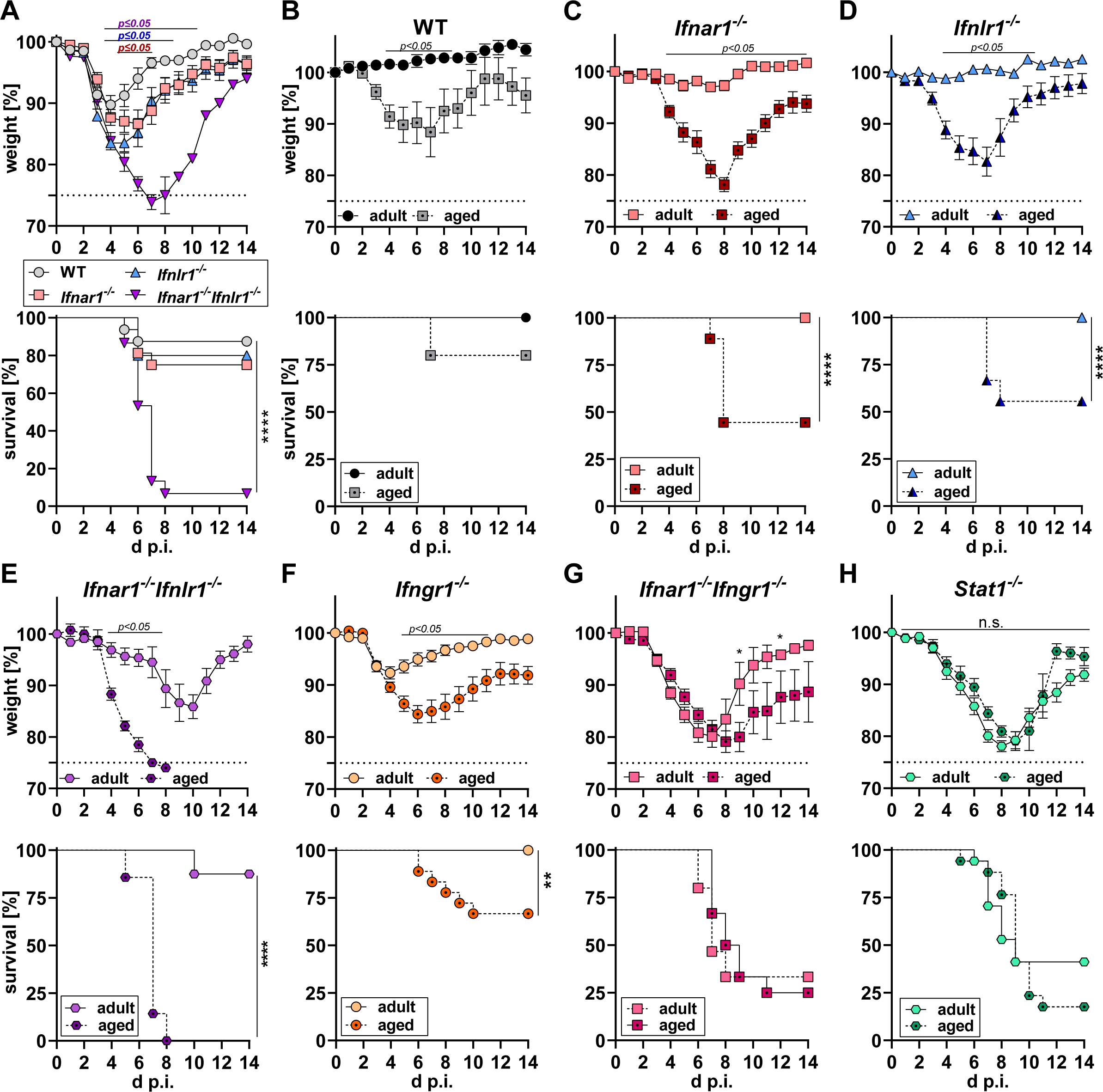
Combined deficiency in type I and type II IFN signaling phenocopies age-dependent disease susceptibility. **A**) Adult (8-20-week-old) WT, *Ifnar1^-/-^*, *Ifnlr1^-/-^*and *Ifnar1^-/-^Ifnlr1^-/-^* mice (n=15-16) were infected with 300 PFU SARS-CoV-2 MA20. Weight loss (upper panel) and survival (lower panel) was monitored for 14 days post infection. Data pooled from two independent experiments. Dashed line indicates experimental endpoint due to animal welfare. Symbols represent mean ± SEM. Weight loss: *p*-values for WT in comparison with *Ifnar1^-/-^*(red), *Ifnlr1^-/-^* (blue) and *Ifnar1^-/-^Ifnlr1^-/-^*(purple) by two-way ANOVA with Tukey’s multiple comparisons test. Survival: \*\*\*\**P*≤0.0001, Log-rank (Mantel-Cox) test. **B-H**) Groups of adult or aged (10-20-week- or 38-60-week-old) mice of the indicated genotypes were infected with 30 PFU SARS-CoV-2 MA20. Weight loss (upper panel) and survival (lower panel) was monitored for 14 days post infection. Dashed line indicates experimental endpoint due to animal welfare. Weight loss: *p*-values calculated by two-way ANOVA with Šídák’s multiple comparisons test. Survival: \**P*≤0.05, \*\*\*\**P*≤0.0001, Log-rank (Mantel-Cox) test. (**B-E**) All mice were infected in parallel and data from a single experiment are shown; n=5-9. (**F**) Data pooled from three independent experiments; n=18-20. (**G**) Data pooled from two independent experiments; n=11-15. (**H**) Data pooled from two independent experiments; n=17.

Collectively, these data suggested that excessive virus replication promoted by an impaired type I IFN system in combination with impaired IFN-γ-mediated immune responses can account for the observed high SARS-CoV-2 disease susceptibility of aged mice.

### Therapeutic administration of IFN-λ prevents SARS-CoV-2-induced lethality and supplementation of IFN-γ reverses the age-dependent disease phenotype

Experiments using *Ifnar1^-/-^* and *Ifnar1^-/-^Ifnlr1^-/-^* mice demonstrated that endogenously produced type III IFNs can partially substitute for a dysfunctional type I IFN immunity^2–4^, thereby protecting against lethal disease progression (**Figure 6A**). Therefore, we evaluated the antiviral potential of IFN-λ as a drug candidate^47^ in the context of a dysfunctional type I IFN immunity. We could demonstrate that a single prophylactic administration of 2 µg IFN-λ1/3^44^ one day prior to infection or a therapeutic regimen of 3 µg/d for one week starting one day after infection, efficiently prevented lethal disease progression in adult *Ifnar1^-/-^* mice (**Figure 7A**). However, modeling advanced age in combination with dysfunctional type I IFN immunity by using aged *Ifnar1^-/-^*mice, we found that a single prophylactic administration of IFN-λ was insufficient to significantly reduce disease burden in such highly susceptible mice (**Figure 7B**), whereas therapeutic administration of IFN-λ showed some residual protective activity by significantly reducing weight loss with a trend towards increased survival rates (**Figure 7B**). Of note, therapeutic application of IFN-λ efficiently protected aged WT animals with functional type I IFN immunity against SARS-CoV-2 induced disease and lethality (**Figure S3B**). Next, we treated aged WT mice daily with 2 µg IFN-γ from -1 to 7 d p. i. to test whether supplementation of IFN-γ could reverse the age-dependent enhanced disease severity upon SARS-CoV-2 infection. Infection with 200 PFU MA20 caused substantial weight loss and lethal disease progression in 5 out of 8 aged WT mice, whereas aged WT mice supplemented with IFN-γ showed a significantly reduced weight loss and an increased survival rate which was nearly identical with the disease course of adult WT mice (**Figure 7C**). Encouraged by the positive results of IFN-λ in adult *Ifnar1^-/-^*and those of IFN-γ in aged WT animals, we next tested a combination of both treatments in an attempt to protect highly susceptible aged mice with a defective type I IFN system. Whereas individual therapeutic treatment regimens with either IFN-λ or IFN-γ in aged *Ifnar1^-/-^* both conferred only limited protection against SARS-CoV-2 induced disease and lethality (**Figure 7B** **and S3C**), the combinatorial therapeutic treatment with IFN-λ1/3 and IFN-γ prevented morbidity exceeding 10 % of body weight loss in an otherwise lethal infection (**Figure 7D**).

**Figure 7.**
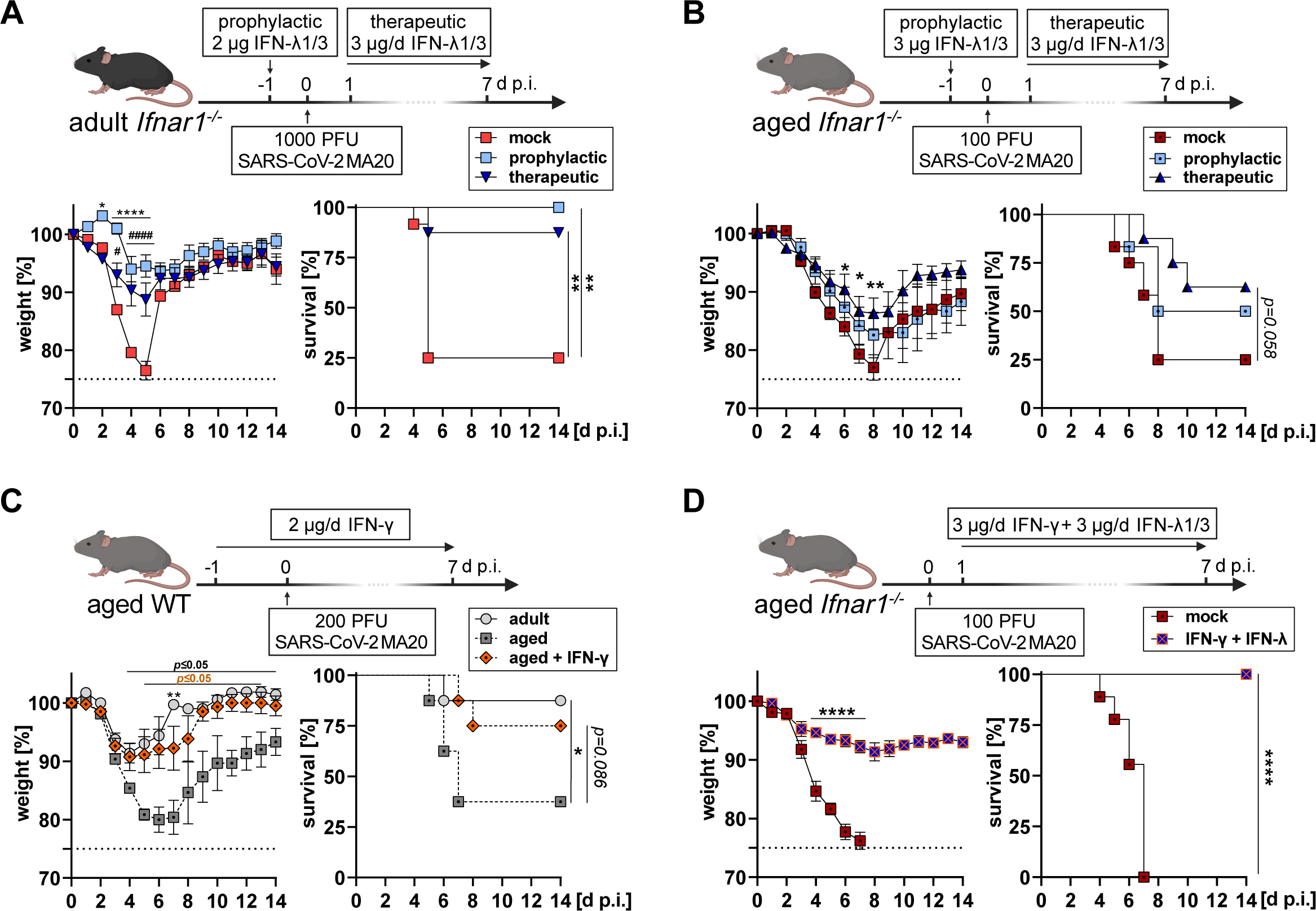
Therapeutic administration of IFN-λ prevents SARS-CoV-2-induced lethality in absence of type I IFN immunity and supplementation of IFN-γ reverses age-dependent disease phenotype. **A)** Groups of adult *Ifnar1^-/-^*mice (9-15-week-old) were either mock-treated (n=12), treated prophylactically by subcutaneous injection of 2 µg IFN-λ1/3 once one day prior to infection (n=6), or treated therapeutically with 3 µg IFN-λ1/3 daily for one week (n=8) starting one day after infection with 1000 PFU SARS-CoV-2 MA20. Dashed line indicates experimental endpoint due to animal welfare. Pooled data from two independent experiments are shown. Symbols represent mean ± SEM. Weight loss: mock in comparison with prophylactic (asterisks) or therapeutic (hashes) by two-way ANOVA with Tukey’s multiple comparisons test; \**P*≤0.05, ^#^*P*≤0.05, \*\*\*\**P*≤0.0001, ^####^*P*≤0.0001. Survival: Log-rank (Mantel-Cox) test; \*\**P*≤0.01. **B)** Groups of aged *Ifnar1^-/-^* (49-60-week-old) mice were either mock-treated (n=12), treated prophylactically by subcutaneous injection of 3 µg IFN-λ1/3 once one day prior to infection (n=6), or treated therapeutically with 3 µg IFN-λ1/3 daily for one week (n=8) starting one day after infection with 100 PFU SARS-CoV-2 MA20. Dashed line indicates experimental endpoint due to animal welfare. Pooled data from two independent experiments are shown. Symbols represent mean ± SEM. Weight loss: mock in comparison with therapeutic by two-way ANOVA with Tukey’s multiple comparisons test; \**P*≤0.05, \*\**P*≤0.01. Survival: Log-rank (Mantel-Cox) test. **C)** Groups of aged WT mice (45-52-week-old) were either mock-treated (n=8) or treated subcutaneously with 2 µg IFN-γ daily (n=8) for nine days starting one day prior to infection with 200 PFU SARS-CoV-2 MA20. Infected but untreated adult WT mice (10-16-week-old; n=8) served as controls. Dashed line indicates experimental endpoint due to animal welfare. Data from a single experiment are shown. Symbols represent mean ± SEM. Weight loss: *P*≤0.05 for mock-treated aged mice in comparison with adult control mice (black) or with IFN-γ-treated aged mice (orange) by two-way ANOVA with Tukey’s multiple comparisons test; \*\**P*≤0.01 comparing adult control mice with IFN-γ-treated aged mice. Survival: Log-rank (Mantel-Cox) test, \**P*≤0.05. **D)** Groups of aged *Ifnar1^-/-^* mice (52-60-week-old) were either mock-treated (n=9) or treated therapeutically with a mixture of 3 µg IFN-λ1/3 and 3 µg IFN-γ daily for one week (n=8), starting one day after infection with 100 PFU SARS-CoV-2 MA20. Dashed line indicates experimental endpoint due to animal welfare. Data from a single experiment are shown. Symbols represent mean ± SEM. Weight loss: mock in comparison with therapeutic treatment group by two-way ANOVA with Šídák’s multiple comparisons test; \*\*\*\**P*≤0.0001. Survival: Log-rank (Mantel-Cox) test; \*\*\*\**P*≤0.0001.

Taken together these data demonstrated that (i) the drug candidate IFN-λ1/3 could efficiently prevent lethal disease progression in adult mice with defective type I IFN immunity, that (ii) supplementation of IFN-γ could reverse the age-dependent enhanced disease progression and that (iii) the combination of both treatments rescued lethally infected aged mice lacking type I IFN responses (**Figure S4**).

## Discussion

Two major risk factors for severe COVID-19 are advanced age^1^ and an impaired IFN-mediated immunity^2–4^. For the rational design of effective therapeutics and prevention strategies targeting the respective risk groups, a better understanding of the disease-causing mechanisms is urgently needed. Small animal models faithfully recapitulating characteristics of human disease are pivotal to overcome these limitations. In this study, we generated a highly pathogenic mouse-adapted SARS-CoV-2 variant that can be used to model mild, moderate or severe COVID-19 as well as the age-associated aggravation of disease in standard inbred mice.

Using this small animal model, we found that the age-dependent increase in disease severity is driven by an impaired interferon response which causes a delayed, insufficient and dysregulated innate and adaptive immune response in the aged host. Transcriptome analyses of infected lungs from mature adult and middle-aged C57BL/6 mice revealed that adult mice initiated a rapid and well-coordinated innate and adaptive immune response, which was associated with high IFN-γ and low IL-10 expression levels. This effective and timely immune response in adults limited viral loads, mediated rapid viral clearance, and efficiently prevented the development of severe disease. In aged mice, by contrast, virus replication was markedly increased which correlated with the absence of effective antiviral immune responses. Instead, aged mice showed strong IL-6- and IL-1-mediated responses associated with low IFN-γ and high IL-10 expression levels. The markedly different IFN-γ to IL-10 ratio likely explains the effective immune response, including NK cell mediated immunity, efficient antigen presentation, lymphocyte activation, and immunoglobulin production in adult mice that leads to the favorable disease outcome compared with aged counterparts (**Figure S4**).

Interestingly, in spite of the strongly increased viral loads and the enhanced disease progression in aged animals, lung tissue damage was comparable between both age groups. This finding possibly indicates a functional impairment of the infected aged lung rather than excessive cellular damage that would be detected using classical histological methods such as H&E stainings. Alternatively, it is possible that aged mice suffered from a systemic manifestation of disease due to overshooting cytokine production^48^ or virus dissemination to other organs. Indeed, RT-qPCR analyses revealed elevated viral RNA levels in both heart and brain samples of aged mice. However, neither signs of inflammation or immune cell infiltrations were detected nor were viral RNA-positive cells found in heart and brain samples from aged mice using highly sensitive RNA in situ hybridization methods, arguing against a systemic dissemination of the virus in aged animals.

To test whether an impaired IFN response would indeed phenocopy the enhanced disease progression in the aged host, we directly compared adult and aged mice with individual or combinatorial deficiencies in the various IFN pathways. We found that type I and type III IFNs limited disease susceptibility and lethality but genetic defects in these pathways did not faithfully recapitulate the age-dependent enhanced disease phenotype. Importantly, however, adult *Ifngr1^-/-^*mice with a deficient type II IFN system suffered from increased disease severity that closely resembled the disease course of aged wild type mice. Nevertheless, aged *Ifngr1^-/-^* mice still showed enhanced disease progression and increased lethality compared with their younger counterparts, indicating that additional mechanisms are promoting the age-dependent disease severity. Indeed, adult *Ifnar1^-/-^Ifngr1^-/-^*mice lacking functional type I and type II IFN signaling, showed the same degree of severe weight loss and poor survival rates as their aged counterparts, demonstrating a defining role of diminished type I and type II IFN responses driving the age-dependent virulence of SARS-CoV-2.

We therefore hypothesize that the age-dependent impairment of IFN-γ-mediated responses causes the impaired NK cell activation, reduced antigen presentation and diminished lymphocyte activation that we see in our transcriptome data, whereas the impaired type I IFN immunity facilitates virus replication due to reduced cell-intrinsic antiviral immunity. In agreement with our finding that adult *Ifngr1*^-/-^ mice phenocopied the disease course of aged wild type animals, we found that treatment of aged mice with IFN-γ prevented severe disease progression, supporting the view that IFN-γ limits age-dependent COVID-19 severity. By comparing children and adults, other researchers found that increased expression levels of PRRs such as MDA5 and RIG-I maintain a tonic IFN-activated antiviral state in the airways which appears to contribute to the high resistance of young individuals against severe COVID-19^16, 17^. However, it is unclear whether such mechanisms, which may explain the elevated baseline of type I and type III IFNs, would affect IFN-γ production. Whereas type I and type III IFNs are produced mainly by dendritic cells as well as infected epithelial cells upon activation of PRRs^11^, IFN-γ is mainly produced by NK and T cells and involves distinct transcription factors such as Eomes and T-bet in its regulation^49^. Of note, adult mice in our study showed significantly increased expression levels of T-bet in their infected lungs compared with aged mice.

Additional studies are required to determine which pathways and cell types are involved in the detection of SARS-CoV-2 infection, and which cells produce the respective types of interferon to prevent severe disease. The identification of cell types that fail to produce IFN-γ and/or fail to adequately respond to this cytokine in the aged organism may enable the development of novel strategies that aim at preventing or reversing such pathological processes. A recent study showed that soluble factors in plasma from severe COVID-19 patients reversibly interfered with the antiviral activity of NK cells and their capacity to mount an appropriate IFN-γ and TNF response that otherwise was associated with a favorable disease outcome^28^. Interestingly, others suggested a potential disease-driving role of IFN-γ in combination with TNF, possibly inducing inflammatory cell death during severe COVID-19 via PANoptosis^50^. However, we observed that the protective effects of endogenous or exogenous IFN-γ during SARS-CoV-2 infection clearly outweigh possible detrimental effects in our infection model. Similarly, two recent studies raised awareness that continuous IFN-λ signaling could hamper with efficient lung repair after viral infection^51, 52^. However, in our COVID-19 mouse model, IFN-λ deficiency led to increased and prolonged viral replication causing enhanced disease and delayed recovery. Vice versa, both prophylactic and therapeutic treatment with IFN-λ strongly reduced viral loads in the lung and efficiently prevented disease deterioration in lethally infected aged mice or adult animals lacking type I IFN immunity. This strongly indicates that the protective effects of IFN-λ during severe SARS-CoV-2 infections outcompete potential adverse effects. Notably, individual therapeutic treatment of infected aged *Ifnar1*^-/-^ mice, a model combining two of the most detrimental risk factors for severe COVID-19 in humans^1, 3^, with either IFN-γ or IFN-λ had limited success in preventing severe disease and lethality. However, the combination of both cytokines proved to be highly efficient. We therefore hypothesize that exogenous IFN-γ rescues the age-related impairment of cellular immune responses, whereas IFN-λ compensates for the compromised cell-intrinsic antiviral immunity in epithelial cells. Both cytokines may also improve DC migration into draining lymph nodes facilitating T cell activation^53^, a process which is impaired in infected aged mice due to elevated levels of prostaglandin D2 and its upstream phospholipase PLA2G2D^8, 54–56^. Interestingly, such elevated PGD2 levels may also drive enhanced disease by interfering with the production of IFN-λ^57^. Both IFN-γ and IFN-λ treatments were already evaluated in various clinical settings. IFN-γ is licensed to treat chronic granulomatous disease^58^ and is discussed as a treatment for idiopathic pulmonary fibrosis^59^. IFN-λ treatment was well tolerated by patients in the context of HCV infections^60–62^, and a recent clinical trial demonstrated a marked efficacy in accelerating viral decline and clearance in SARS-CoV-2 infected patients^24^. Given the continuous burden of the currently ongoing and the threat of future pandemics, preparations should be taken to have broad-spectrum antivirals like type II and type III interferons accessible for the highly exposed and vulnerable if needed.

## Material and Methods

### Viruses and cell lines

The virus strain SARS-CoV-2 MA containing the amino acid changes Q498Y and P499T in the spike protein^37^ has been generated by transformation-associated recombination in yeast as previously described^63^. Briefly, the mutations resulting in the Q498Y and P499T amino acid changes in the spike protein were generated in plasmids containing fragments 9 and 10 by using primers mSARSv3Fg9+10-F (5’-ATATGGTTTCtacacgACTAATGGTGTTGGTTAC) and mSARSv3Fg9+10-R (5’-GATTGTAAAGGAAAGTAACAATTAAAAC) by Q5 site-directed mutagenesis (New England Biolabs). Passage one SARS-CoV-2 MA stocks produced using Vero E6 cells were used in experiments. The identity of the resulting recombinant SARS-CoV-2 MA was confirmed by sequencing.

Plaque-purified virus stocks from passage 20 lung homogenates of passaging series DKO A and B were grown using Vero E6 cells. Plaque-purified virus stock derived from passaging series DKO A was plaque-purified a second time using Vero E6 cells to generate a SARS-CoV-2 MA20 virus stock for experiments. Briefly, virus stocks were produced by inoculating confluent Vero E6 cells with virus diluted in Opti-MEM (Gibco) 0.3 % BSA (Sigma-Aldrich) for 2 h at 37 °C with 5 % CO_2_ after removal of cell culture medium (DMEM (Gibco) containing 10 % FCS (Anprotec), 20 U/ml penicillin, and 20 µg/mL streptomycin (Gibco)) and washing with PBS. After removal of infection medium, cells were washed with PBS and DMEM containing 2 % FCS and 20 mM HEPES (Roth) was added. After incubation for 48-72 h at 37 °C with 5 % CO_2_, cell culture supernatants were harvested, cleared by centrifugation, aliquoted and stored at -80 °C until further use. Viral load in virus stocks was determined by plaque assay using Vero E6 cells. Purity of virus stocks was confirmed by next-generation sequencing. Sequenced B.1 (Muc-IMB-1/2020) and B.1.617.2 (Delta) virus stocks were provided by Georg Kochs^64^.

African green monkey kidney Vero E6 cell line (ATCC CRL-1586) and human lung adenocarcinoma Calu-3 cell line (ATCC HTB-55) were purchased from ATCC. All virus infection experiments were performed under BSL-3 conditions.

### Mice

C57BL/6JRj, BALB/cJRj and 129S2/SvPasOrlRj mice were purchased from Janvier Labs. B6.A2G-*Mx1,* B6.A2G-*Mx1-Ifnar1^−/−^*, B6.A2G-*Mx1-Ifnlr1^−/−^*, B6.AG2*-Mx1-Ifnar1^−/−^Ifnlr1^−/−^*, B6.*Ifngr1^-/-^,* B6.*Ifnar1^-/-^Ifngr1^-/-^,* and B6.A2G-*Mx1*-*Stat1^-/-^* were bred and kept at the animal facilities of the University Medical Center Freiburg.

Mature adult 8- to 20-week-old mice, designated “adult”, and middle-aged 36- to 60-week-old mice, designated “aged”, were used in experiments. Animals of both sexes were used. All experimental groups were sex- and age-matched.

### Infection of mice and IFN treatments

Mice were anaesthetized using 1.8-2.8 Vol % isoflurane in O2 and subsequently inoculated with 40 µl PBS 0.1 % BSA containing the indicated dose of the respective virus variant via their nostrils. Infected mice were monitored daily for weight loss and clinical signs of disease for at least two weeks. Experimental end points when mice were euthanized by cervical dislocation were defined as ≥25 % weight loss or ≥20 % weight loss for longer than two days. To collect organ samples mice were euthanized by cervical dislocation. For histopathological analyses, mice were anaesthetized using ketamine/xylazine and fixed by transcardial perfusion with 10 % formalin solution and stored in 10 % formalin solution at 4 °C until further processing.

To determine viral loads by plaque assay, organs were homogenized 3-4 times in 800 µl PBS at 6.5 m/s for 18 s using a FastPrep™ tissue homogenizer (MP Biomedicals). Homogenates were centrifuged at 10 000 rpm for 10 min, supernatants collected and stored at -80 °C until further processing.

Nasal swabs were collected using a wet cotton-swab and stored in 650 µl Opti-MEM 0.3 % BSA at -80 °C. Samples were thawed at 20 °C with 1 400 rpm for 15 min on a thermoshaker and viral load determined by plaque assay.

For intranasal treatment, isoflurane-anaesthetized mice were mock-treated with PBS 0.1 % BSA or inoculated with the respective doses of IFN-α ^65^ or human IFN-λ1/3^44^ in a 40 µl volume via their nostrils.

For subcutaneous treatment, mice were mock-treated with PBS 0.1 % BSA or injected with the respective doses of human IFN-λ1/3, mouse IFN-γ or a combination of both in a 100 µl volume using a 30 G syringe.

### Plaque assay

Ten-fold serial dilutions of infectious samples were prepared in Opti-MEM 0.3 % BSA and added to PBS-washed confluent Vero E6 cells, incubated for 2 h at 37 °C with 5 % CO_2_ before inoculum was removed, and cells overlaid with DMEM containing 0.1 % BSA, 1.5 % Avicel (FMC BioPolymer), 0.5 mg/ml L-glutamine (Roth), 20 mM HEPES 20 U/ml penicillin, and 20 µg/mL streptomycin and incubated for 72 h at 37 °C with 5 % CO_2_. After removal of Avicel medium, cells were fixed using 10 % formalin solution and stained with 1 % crystal violet in H2O containing 20 % ethanol, each step for at least 25 min. Plaques were counted and used to calculate virus titers defined as PFU per ml.

To plaque-purify clonal isolates, cells infected with serial dilutions were overlaid with DMEM containing 2 % FCS, 0.6 % agar (Oxoid Germany), 0.01 % DEAE Dextran (Sigma), and 0.1 % NaHCO3 (Merck) and plaques picked 48-72 h p. i. using a thin filter-tip containing 2 µl PBS.

### Cell culture, virus growth curves & plaque reduction neutralization test

Vero E6 and Calu-3 cells were cultured in DMEM and MEM, respectively, containing 10 % FCS, 20 units/ml penicillin and 20 µg/mL streptomycin.

For virus growth curves, Vero E6 or Calu-3 cells seeded in 24-well plates were washed with PBS and then infected with the respective virus strain by incubating the cells with virus containing Opti-MEM 0.3 % BSA for 2 h at 37 °C with 5 % CO_2_ using an MOI of 0.001. Cells were washed with PBS and DMEM 2 % FCS added. Cell culture supernatants were collected at the indicated time points and viral loads determined by plaque assay on Vero E6 cells.

Serological neutralization tests were performed using sera from vaccinated individuals 10- to 25-weeks post-vaccination with the second dose of Comirnaty (BioNTech/Pfizer) or Spikevax (Moderna). Serial 2-fold dilutions of sera in Opti-MEM 0.3 % BSA were incubated for 1 h with 100 PFU of the indicated SARS-CoV-2 variant. The mixture was then added to Vero E6 cells and incubated for 1.5 h at 37 °C with 5 % CO_2_. After removal of the inoculum, cells were overlaid with DMEM containing 2 % FCS, 0.6 % agar, 0.01 % DEAE Dextran, and 0.1 % NaHCO3 and incubated for 72 h at 37 °C with 5 % CO_2_. Cells were fixed for 20 min using 10 % formalin solution, stained for 20 min with 1 % crystal violet in H2O containing 20 % ethanol and plaques were counted. Plaque reduction was calculated relative to mock-treated controls. 50 % neutralizing titers (NT50) were calculated by non-linear fit least squares regression (constraints: 0 and 100).

### RNA isolation and RT-qPCR

For RNA isolation, organ samples were homogenized four times in 800 µl TRI Reagent® (Zymo Research Corporation) at 6.5 m/s for 20 s using a FastPrep™ tissue homogenizer (MP Biomedicals). Samples were centrifuged at 10 000 rpm for 5 min and supernatant diluted in TRI Reagent® (1:2 to 1:32) was used for RNA extraction using the Direct-zol™ RNA Miniprep kit according to the manufacturer’s instructions (Zymo Research Corporation).

cDNA was reverse-transcribed from 750 ng total RNA per sample using the LunaScript® RT SuperMix kit (New England Biolabs) and served as template for amplification of genes of interest (*Ubc*, QuantiTect® Primer Assay, Cat. No. QT00245189; *Tnf*, QuantiTect® Primer Assay, Cat. No. QT00104006; *Ifna4*, QuantiTect® Primer Assay, Cat. No. QT01774353; *Ifnl2/3*, Applied Biosystems, Cat. No. mm0420156_gH; *Ifnb1*, forward: 5’-CCTGGAGCAGCTGAATGGAA-3’, reverse: 5’-CACTGTCTGCTGGTGGAGTTCATC-3’, probe: 5’-[6FAM]CCTACAGGGCGGACTTCAAG[BHQ1]−3’; *Isg15*, forward: 5’-GAGCTAGAGCCTGCAGCAAT-3’, reverse: 5’-TTCTGGGCAATCTGCTTCTT-3’; *Stat1*, forward: 5’-TCACAGTGGTTCGAGCTTCAG-3’, reverse: 5’-CGAGACATCATAGGCAGCGTG-3’; *Mx1*, forward: 5’-TCTGAGGAGAGCCAGACGAT-3’, reverse: 5’-ACTCTGGTCCCCAATGACAG-3’; *Il6*, forward: 5’-TCGGAGGCTTAATTACACATGTTCT-3’, reverse: 5’-GCATCATCGTTGTTCATACAATCA-3’) using SYBR™ Green PCR Master Mix (Applied Biosystems) or TaqMan® Universal PCR Master Mix (Qiagen) and a QuantStudio™ 5 Real-Time PCR System (Applied Biosystems). The increase in mRNA expression was determined by the 2^-ΔCt^ method relative to the expression of the indicated housekeeping gene.

Viral RNA was quantified by one-step RT-qPCR using the AgPath-ID™ One-Step RT-PCR (Applied Biosystems) reagents and the 2019-nCoV RT-qPCR primers (E_Sarbeco)^66^ specific for the viral E gene. Serial dilutions of a defined RNA standard was used for absolute quantification as previously described^67^.

### Virus genome sequencing

RNA was extracted from 50 µl lung homogenate supernatant using the NucleoSpin^®^ RNA Mini Kit according to the manufacturer’s protocol (MACHEREY-NAGEL). cDNA was reverse-transcribed from extracted RNA using random hexamer primers and Superscript III (ThermoFisher) followed by PCR tiling of the entire SARS-CoV-2 genome (ARTIC V3 primer sets; https://github.com/artic-network/artic-ncov2019) producing ∼400 bp long overlapping amplicons that were used to prepare the sequencing library. Amplicons were purified with AMPure magnetic beads (Beckman Coulter) and QIAseq FX DNA Library Kit (Qiagen) used to prepare indexed paired-end libraries for Illumina sequencing. Normalized and pooled sequencing libraries were denatured with 0.2 M NaOH. Libraries were sequenced on an Illumina MiSeq using the 300-cycle MiSeq Reagent Kit v2.

De-multiplexed raw reads were subjected to a customized Galaxy pipeline based on bioinformatics pipelines on usegalaxy.eu^68^. Raw reads were pre-processed with fastp v.0.20.1^69^ and mapped to the SARS-CoV-2 Wuhan-Hu-1 reference genome (Genbank: NC_045512) using BWA-MEM v.0.7.17^70^. Primer sequences were trimmed using ivar trim v1.9 (https://andersen-lab.github.io/ivar/html/manualpage.html). Variants (SNPs and INDELs) were called using the ultrasensitive variant caller LoFreq v2.1.5^71^, demanding a minimum base quality of 30 and a coverage ≥ 10-fold. Called variants were filtered based on a minimum variant frequency of 10 %. Effects of mutations were automatically annotated in vcf files using SnpEff v.4.3.1^72^. Consensus sequences were constructed using bcftools v.1.1.0^73^. Regions with low coverage (>20-fold) or variant frequencies between 30 and 70 % were masked with N.

A customized R script was used to plot variant frequencies that were detected by LoFreq as a heatmap (github.com/jonas-fuchs/SARS-CoV-2-analyses) which is also available on usegalaxy.eu (“Variant Frequency Plot”).

### RNA-sequencing

RNA was isolated from infected lungs as described above. RNA-sequencing was performed on the HiSeq 4000 system (Illumina) with Single End 75 bp reads. Read quality trimming and adaptor removal was carried out using Trimmomatic (version 0.36). The nf-core/rnaseq pipeline (version 3.0;^74^) written in the Nextflow domain specific language (version 19.10.0;^75^) was used to perform the primary analysis of the samples in conjunction with Singularity (version 2.6.0;^76^). All data was processed relative to the mouse GRCm38 genome downloaded from Ensembl. Gene counts per gene per sample were obtained using the RSEM-STAR (^77, 78^) option of the pipeline and they were imported on DESeq (v1.28.0, ^79^) within R environment v4.0.2 for differential expression analysis. Gene Ontology and Gene Set Enrichment analysis (GSEA) were carried out using R package Cluster Profiler (v3.16). For GSEA, gene lists ranked using the Wald statistic were used. Pre-ranked analyses were carried out using C5 ontology GO biological process gene sets from the Molecular Signatures database (MSigDB, v7.2). Gene signatures were considered significant if FDR *q*-value ≤ 0.05. ggplot2, RColorBrewer, ComplexHeatmap were used for plotting purposes. Ingenuity Pathway Analysis was performed using differentially expressed genes (fold change ≥ 1.5, *p*adj ≤ 0.05). The indicated -log(*p*-values) were calculated using the Benjamini-Hochberg method of multiple testing correction.

### Histopathology, immunohistochemistry, RNA in-situ-hybridization and scoring

Mice, after cardiac perfusion as described above, were immersion-fixed with 10 % neutral-buffered formalin solution. Tissue was embedded in paraffin, including the whole lung, 3-4 trimmed sections of the decalcified nasal cavity, a sagittal section of the brain, and a longitudinal section of the heart. Samples were cut in 2-3-μm-thick sections and stained with hematoxylin and eosin (HE) for light microscopical examination. Consecutive slides were processed for immunohistochemistry (IHC). A polyclonal serum detecting the nucleocapsid protein of SARS-CoV-2 (Rockland Immunochemicals # 200-401-A50, PA, USA) was used according to standardized procedures of the avidin-biotin-peroxidase complex-method (ABC, Vectastain Elite ABC Kit, Burlingame, CA, USA). Briefly, 2-3 µm sections were mounted on adhesive glass slides, dewaxed in xylene, followed by rehydration in descending graded alcohols. Endogenous peroxidase was quenched with 3 % hydrogen peroxide in distilled water for 10 minutes at room temperature. Antigen heat retrieval was performed in 10 mM citrate buffer (pH 6) for 20 minutes in a pressure cooker. Nonspecific antibody binding was blocked for 30 minutes at room temperature using normal goat serum, diluted 1:2 in PBS. The primary polyclonal serum was applied overnight at 4 °C (1:3000, diluted in TRIS buffer), the secondary biotinylated goat anti-rabbit antibody was applied for 30 minutes at room temperature (Vector Laboratories, Burlingame, CA, USA, 1:200). Color was developed by incubating the slides with freshly prepared avidin-biotin-peroxidase complex (ABC) solution (Vectastain Elite ABC Kit; Vector Laboratories), followed by exposure to 3-amino-9-ethylcarbazole substrate (AEC, Dako, Carpinteria, CA, USA). Sections were counterstained with Mayer’s haematoxylin and coverslipped. As negative control, a consecutive section was labelled with an irrelevant antibody detecting M protein of Influenza A virus (ATCC clone HB-64) and a positive control slide was included in each run.

To validate RT-qPCR data, selected tissues (heart and brain) were tested with RNA in situ hybridization (RNA ISH). The RNAScope 2-5 HD Reagent Kit-Red (ACD, Advanced Cell

Diagnostics, Newark, CA) was used as previously published^67^. For hybridization, RNAScope® probes were custom-designed by ACD for SARS-CoV-2 nucleocapsid. The specificity of the probes was verified using a positive control probe detecting RNA encoding for peptidylprolyl isomerase B (cyclophilin B, ppib) and a negative control probe detecting RNA encoding for dihydrodipicolinate reductase (DapB). In addition, to identify subtle inflammation in brain and heart samples from infected 10-week (n=10) and 40-week-old (n=6) C57BL/6, hearts were evaluated for the presence of CD3-positive T cell infiltrates and brains evaluated for the presence of CD3-positive T cells and Iba-1-positive microglial cells/macrophages as described in ^80^.

All sides were scanned using a Hamamatsu S60 scanner and evaluation was done using NDPview.2 plus software (Version 2.8.24, Hamamatsu Photonics, K.K. Japan). The left lung lobe was evaluated for histological changes based on HE staining applying criteria given in **Supplementary Table S1**.

Following IHC, viral antigen was semi-quantitatively recorded in the nasal cavity on ordinal scores using the tiers 0 = no antigen, score 1 = up to 3 foci, score 2 = more than 3 distinct foci, score 3 = coalescing foci, score 4 = >80 % antigen positive. The left lung lobe was evaluated using a 200 x 200 µm grid, positive grids were recorded, and the percentage of positive grids was calculated. Target cells were identified based on the cellular phenotype and their location as bronchial epithelium, alveolar macrophages, and alveolar epithelium, including type 1 and type 2 pneumocytes. For heart and brain samples, IHC-based antigen labeling as well as RNA ISH-based genome detection was recorded as present or absent. Evaluation and interpretation were performed by a board-certified pathologist (DiplECVP) in a masked fashion using the post-examination masking method^81^.

### Ethics and biosafety

The generation of recombinant SARS-CoV-2 MA was approved by the Swiss Federal Office for Public Health (permission A202819).

All work performed at the University Medical Center Freiburg concerning virus isolation, cell culture, and mouse infection experiments with infectious SARS-CoV-2 viruses was conducted in Biosafety Level 3 (BSL-3) laboratories at the Institute of Virology, Freiburg, as approved by the Regierungspräsidium Tübingen (#UNI.FRK.05.16-29). All animal work conducted at the University Medical Center Freiburg and the Francis Crick Institute followed the German animal protection law, or the Animals (Scientific Prpcedures) Act 1986, respectively, and was approved by the respective local animal welfare committee (Regierungspräsidium Freiburg #35-9185.81/G-20/91) or the UK Home Office London (Project Licence No. P9C468066). Animal infection experiments were performed consistent with procedures of the Federation for Laboratory Animal Science Associations and the national animal welfare body.

### Human samples

Sera were obtained from eight vaccinees 10- to 25-weeks after a second dose of the Comirnaty (BioN-TECH/Pfizer) or Spikevax (Moderna) vaccine. Written informed consent was obtained from participants and the study was conducted according to federal guidelines and local ethics committee regulations (Albert-Ludwigs-Universität, Freiburg, Germany: No. F-2020-09-03-160428 and no. 322/20).

### Material, data and code Availability

Material and reagents generated in this study will be made available upon installment of a Material Transfer Agreement (MTA).

Genomic sequence of SARS-CoV-2 MA20, which was generated in this study, has been deposited to _______. Passaging and plaque-purified deep sequencing data have been uploaded to _______. RNA sequencing data are available in GEO under accession code GSE190674.

### Statistical analyses

Data visualization and analyses were performed using GraphPad Prism 9.0 and R version 3.5.1. Specific statistical tests, numbers of animals and / or replicates, and further definitions of precision measures can be found in the respective figure legends or method details. *P*-values are indicated in figures or figure legends.

## Acknowledgements

We thank P. Staeheli for advice and constructive comments, and the animal-care technicians of the University Medical Center Freiburg for their excellent work and support. This work was supported by the Bundesministerium fuer Bildung und Forschung (BMBF) through the Deutsches Zentrum fuer Luft- und Raumfahrt, Germany, (DLR, grant number 01KI2077) and by the Federal State of Baden-Wuerttemberg, Germany, MWK-Sonderfoerdermaßnahme COVID-19/AZ.:33-7533.-6-21/7/2 to MS, Deutsche Forschungsgemeinschaft (DFG), Project no. 453012513 to MB, and by the Swiss National Science Foundation SNSF as a part of NCCR RNA&Disease, a National Centre of Competence (or Excellence) in Research (grant number 182880) and the German Research Foundation (DFG, SPP1596) to VT. The work of SC, ML and AW was funded by the Francis Crick Institute, which receives its core funding from Cancer Research UK (FC001206), the UK Medical Research Council (FC001206), and the Wellcome Trust (FC001206). For the purpose of Open Access, the author has applied a CC BY public copyright licence to any Author Accepted Manuscript version arising from this submission. The funders had no role in the study design, data analysis, data interpretation, and in the writing of this report. All authors had full access to the data in the study and accept responsibility to submit for publication. The authors declare no competing interests. Illustrations were created with BioRender.com.

---

**Figure S1.**
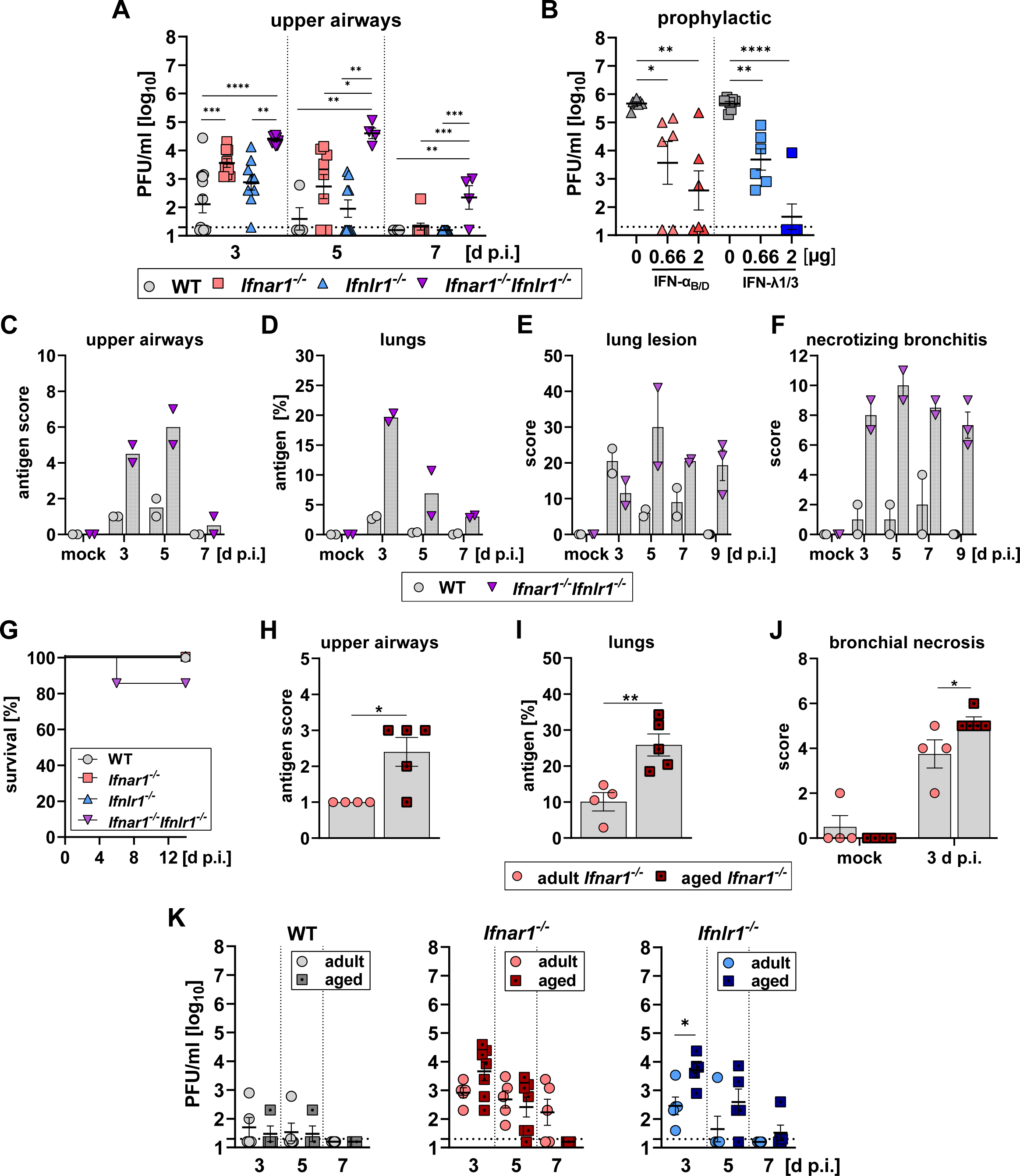
Increased and prolonged replication of SARS-CoV-2 MA in mice lacking type I and/or type III IFN receptors. **A)** Groups of adult mice (8-18-week-old) of the indicated genotypes were infected with 10^5^ PFU SARS-CoV-2 MA. Upper airways were harvested at the indicated time points and viral load determined by plaque assay on Vero E6 cells. Data pooled from five independent experiments. Symbols represent individual mice (n=4-13 per group) and bars indicate mean ± SEM. Dashed line indicates detection limit. \**P*≤0.05, \*\**P*≤0.01, \*\*\**P*≤0.001, \*\*\*\**P*≤0.0001, one-way ANOVA with Tukey’s multiple comparisons test. **B)** Groups of 16-24 week-old *Ifnlr1*^-/-^ (triangles) and *Ifnar1*^-/-^ mice (squares) were intranasally treated with the indicated dose of IFN-α_B/D_ or IFN-λ1/3, respectively, or mock-treated one day prior to infection with 10^5^ PFU SARS-CoV-2 MA. Lung viral loads on day 3 p. i. were determined by plaque assay on Vero E6 cells. Data from a single experiment are shown. Symbols represent individual mice (n=6-7 per group) and bars indicate mean ± SEM. Dashed line indicates detection limit. \*\*\*\**P*≤0.0001, one-way ANOVA with Tukey’s multiple comparisons test. **C-F)** Groups of adult (8-10-week-old) WT or *Ifnar1^-/-^ Ifnlr1^-/-^* mice were mock-treated or infected with 10^5^ PFU SARS-CoV-2 MA (n=2-3 per group) and prepared for histological analyses by cardiac perfusion. Antigen and histopathologic lesion scores for lungs and necrotizing bronchitis were quantified as described in materials and methods section. Data pooled from two independent experiments are shown. Symbols represent individual mice and bars indicate mean ± SEM. **G)** Survival graph corresponding to Figure 1D. **H-J)** Groups of adult (8-10-week-old; n=4) or aged *Ifnar1*^-/-^ mice (36-52-week-old; n=5) were infected with 10^5^ PFU SARS-CoV-2 MA. Mice were prepared for histological analyses by cardiac perfusion on day three post infection. Antigen and histopathologic lesion scores for bronchial necrosis were quantified as described in materials and methods section. \**P*≤0.05, \*\**P*≤0.01, unpaired t test. **K**) Groups of adult or aged mice (8-12-week- or 40-60-week-old; n=4-7) of the indicated genotypes were infected with 10^5^ PFU SARS-CoV-2 MA. Upper airways were harvested at the indicated time points and viral loads determined by plaque assay on Vero E6 cells. Data pooled from four independent experiments are shown. Symbols represent individual mice and bars indicate mean ± SEM. Dashed line indicates detection limit. \**P*≤0.05, unpaired t test.

**Figure S2.**
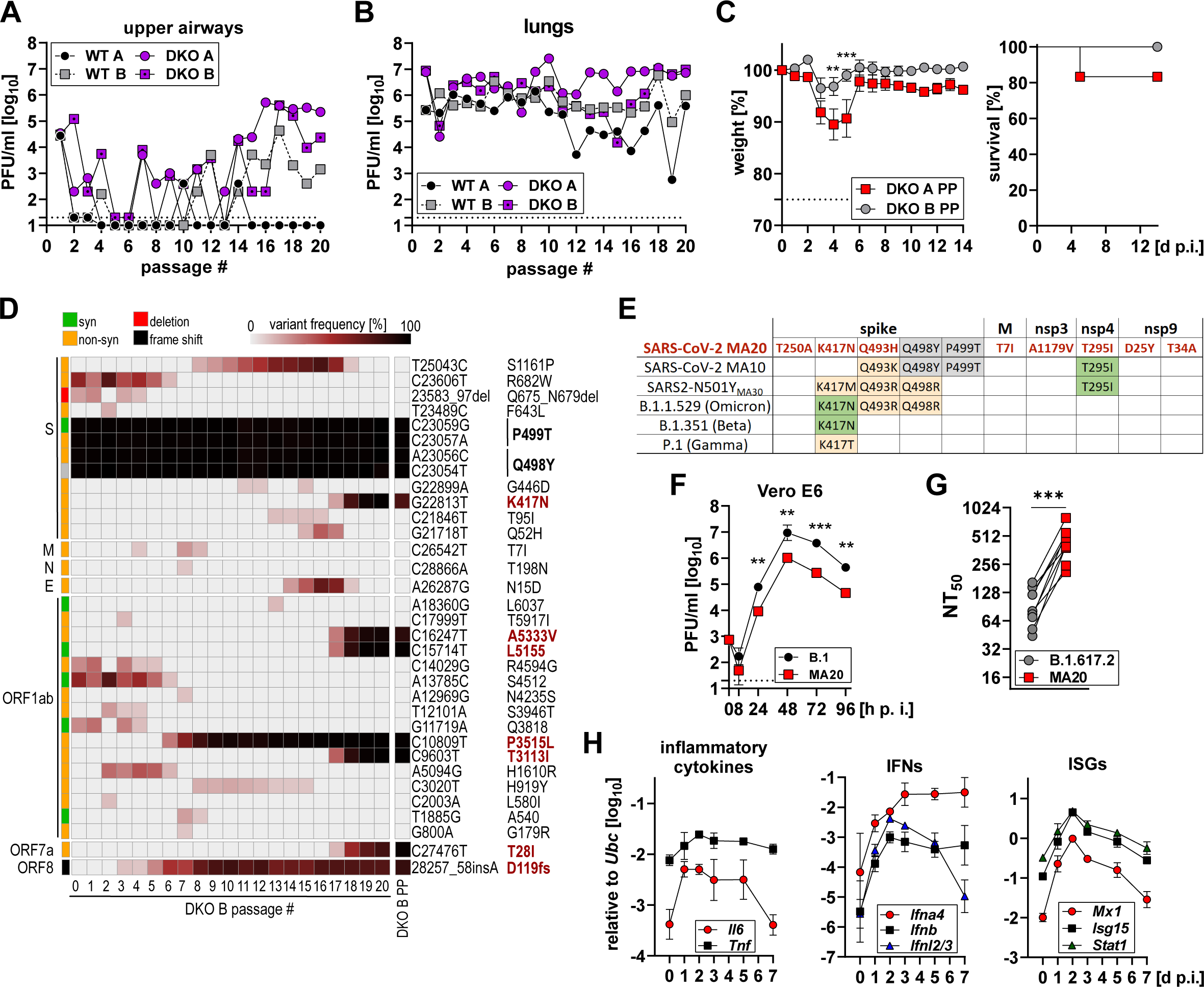
Serial passaging of MA SARS-CoV-2 in IFN receptor-deficient C57BL/6 mice allows for rapid host adaptation. **A-B)** Viral load on day 3 p. i. in upper airways (**A**) and lungs (**B**) for passaging series WT A, WT B, DKO A and DKO B determined by plaque assay on Vero E6 cells. Dashed line indicates detection limit. **C)** Weight loss (left panel) and survival (right panel) of adult C57BL/6 WT mice (13-15-week-old; n=6 per group) infected with 5x10^3^ PFU of plaque-purified virus stocks DKO A PP and DKO B PP. Data from a single experiment are shown. Symbols represent individual mice and bars indicate mean ± SEM. Dashed line indicates experimental endpoint due to animal welfare. \*\**P*≤0.01, \*\*\**P*≤0.001, two-way ANOVA with Šídák’s multiple comparisons test. **D)** Variant frequency plot from next-generation sequencing results for the passaging series DKO B and the plaque-purified “DKO B PP” virus stock. Variant frequencies are shown in comparison to Wuhan-Hu-1 (NC_045512.2). Amino acid changes present in DKO B PP are indicated in bold, changes in comparison to SARS-CoV-2 MA are highlighted in red. **E)** Table indicating similar (orange) or identical (green) amino acid changes present in SARS-CoV-2 MA10, SARS2-N501_MA30_, B1-1-529 (Omicron), B.1.351 (Beta) and P.1 (Gamma) in comparison with amino acid changes present in SARS-CoV-2 MA20 (red). Reference sequence: Wuhan-Hu-1 (NC_045512.2). Amino acid changes highlighted in grey were already present in SARS-CoV-2 MA. **F)** Comparative growth curves of B.1 and SARS-CoV-2 MA20 on VeroE6 cells infected with an MOI of 0.001. Virus replication was quantified by plaque assay on Vero E6 cells. Data from a single experiment performed in duplicates are shown. Dashed line indicates detection limit. \*\**P*≤0.01, two-way ANOVA with Tukey’s multiple comparisons test. **G)** Comparative neutralization by plaque reduction neutralization test of SARS-CoV-2 MA20 and B.1.617.2 (Delta) using sera from vaccinated individuals. Symbols represent mean value for each individual determined in three independent assays. \*\*\**P*≤0.001, paired t test. **H)** Adult C57BL/6 mice (10-week-old; n=5 per group) were infected with 10^3^ PFU of SARS-CoV-2 MA20. Upper airways were harvested at the indicated time points and gene expression levels of *Il6*, *Tnf*, *Ifna4*, *Ifnb*, *Ifnl2/3*, *Mx1*, *Isg15* and *Stat1* were determined relative to *Ubc* by RT-qPCR. Symbols represent mean ± SD.

**Figure S3.**
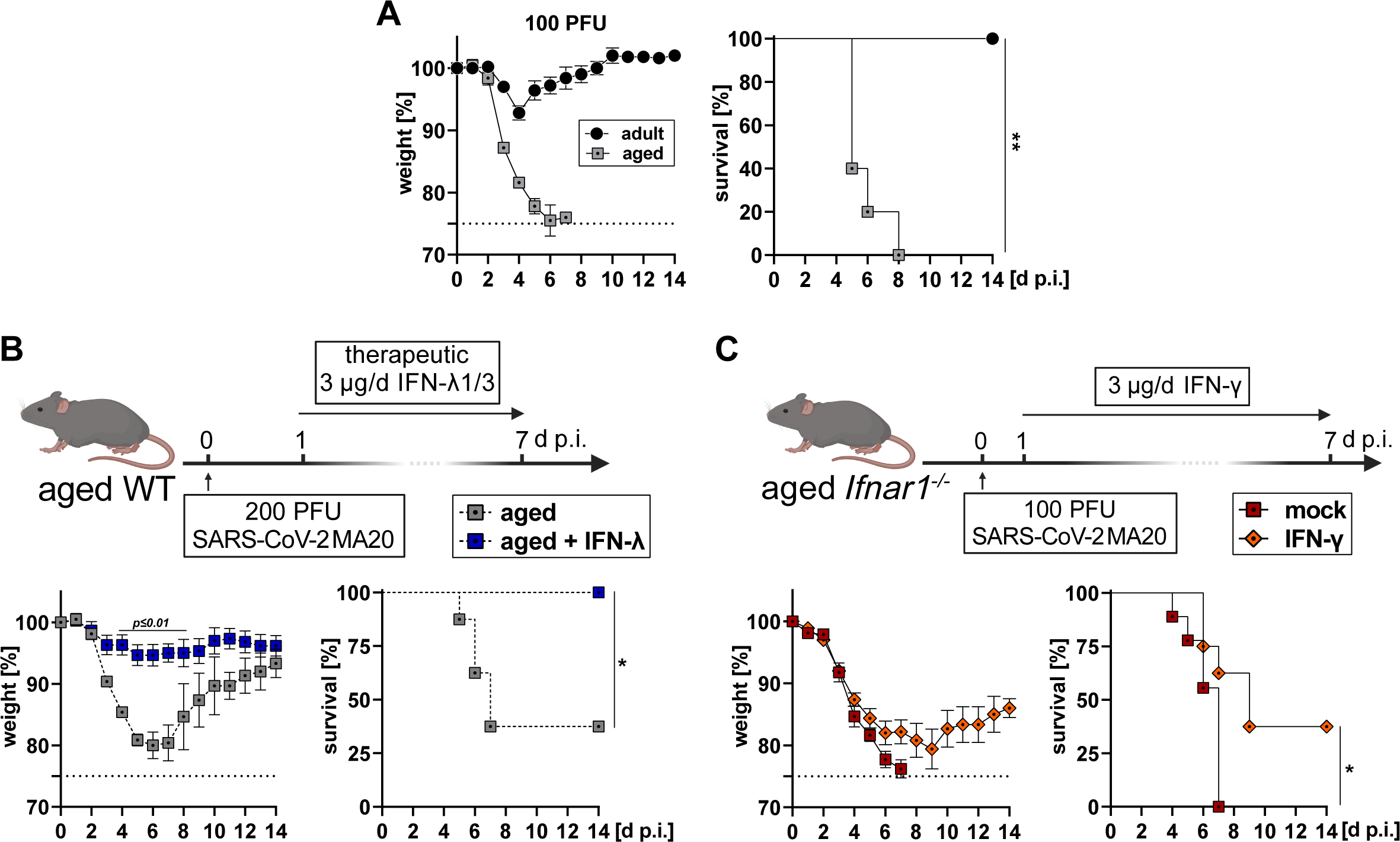
Therapeutic administration of IFN-λ or IFN-γ reduces SARS-CoV-2 induced lethality in aged mice. **A)** Groups of adult or aged C57BL/6 (10-week- or 40-week-old; n = 5 per group) were infected with 100 PFU SARS-CoV-2 MA20. Weight loss (left panel) and survival (right panel) were monitored for 14 days post infection. Data from a single experiment are shown. Symbols represent mean ± SEM. Dashed line indicates experimental endpoint due to animal welfare. Survival: \*\**P*≤0.01, Log-rank (Mantel-Cox) test. Dataset for infected 10-week-old mice is also blotted in Figure 2G. **B)** Groups of aged WT mice (45-56-week-old) were either mock-treated (n=8) or treated therapeutically with 3 µg IFN-λ1/3 (n=8) daily for one week starting one day after infection with 200 PFU SARS-CoV-2 MA20. Mock-treated control group is the same as depicted in Figure 7C. Data from a single experiment are shown. Dashed line indicates experimental endpoint due to animal welfare. Symbols represent mean ± SEM. Weight loss: *P*≤0.05 by two-way ANOVA with Šídák’s multiple comparisons test. Survival: Log-rank (Mantel-Cox) test, \**P*≤0.05. **C)** Groups of aged *Ifnar1^-/-^* mice (52-60-week-old) were either mock-treated (n=9) or treated therapeutically by subcutaneous injection of 3 µg IFN-γ daily for one week (n=8) starting one day after infection with 100 PFU SARS-CoV-2 MA20. Mock-treated control group is the same as depicted in Figure 6D. Data from a single experiment are shown. Dashed line indicates experimental endpoint due to animal welfare. Survival: Log-rank (Mantel-Cox) test, \**P*≤0.05.

**Figure S4.**
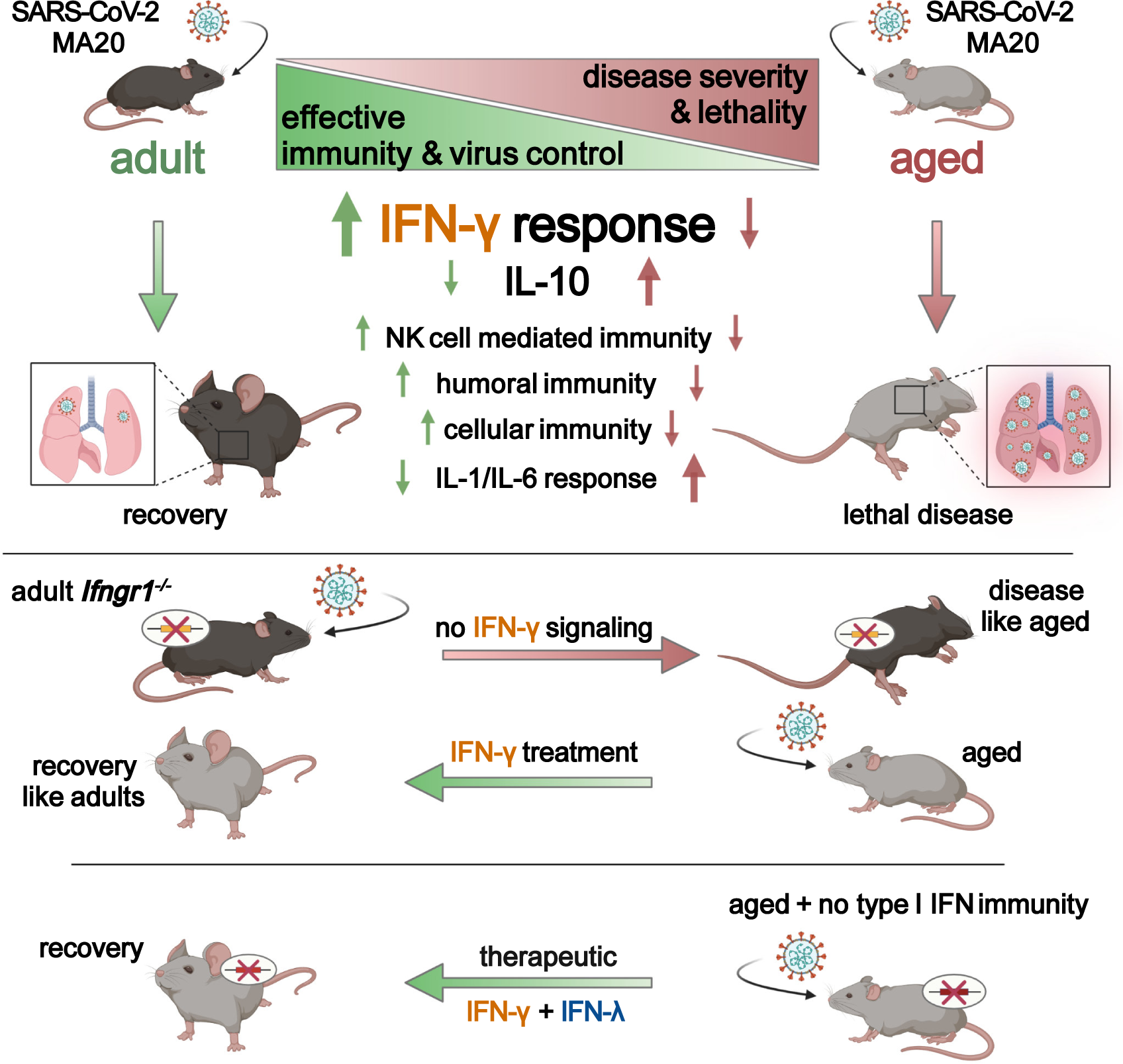
Impaired immune response drives age-dependent virulence of SARS-CoV-2. Graphical summary illustrating the age-dependent impairment of immune responses and suggested intervention strategies.

**Supplementary Table S1:** Histological scoring criteria

| Criteria no. | Criteria | Scoring description* |
| --- | --- | --- |
| 1 | atelectasis (pneumonia-associated), area affected | 0 = no change, 1 = focal to oligofocal <5%, 2 = multifocal (6-40%), 3 = coalescing (41-80%), 4 = diffuse >80% |
| 2 | infiltrates alveolar, area affected | 0 = no change, 1 = focal to oligofocal <5%, 2 = multifocal (6-40%), 3 = coalescing (41-80%), 4 = diffuse >80% |
| 3 | infiltrates alveolar, grade | 0 = no infiltrates, 1 = 1-2 cells, 2 = 2-3 cells, 3 = almost filling, 4 = full atelectasis, grade is representative for the lobe |
|  | description: predominant cell type | granulocytes, lymphocytes, macrophages, plasma cells, mixture (no predominant cell type) |
| 4 | infiltrates interstitial, area affected | 0 = no change, 1 = focal to oligofocal <5%, 2 = multifocal (6-40%), 3 = coalescing (41-80%), 4 = diffuse >80% |
| 5 | infiltrates interstitial, grade | 0 = no infiltrates, 1 = subtle, 2 = thickened septa but regular alveolar pattern, 3 = thickened septa partially obscured alveolar architecture, 4 = full atelectasis; grade is representative for lobe |
|  | description: predominant cell type | granulocytes, lymphocytes, macrophages, plasma cells, mixture (no predominant cell type) |
| 6 | infiltrates peribronchial (incl glands), occurrence | 0 = not present, 1 = present in 1-3 foci, 3 = significantly present in >3 foci |
| 7 | infiltrates peribronchial, grade | = cell layers; 1=1, 2=2-3, 3=4-5, 4≥6 give max grade |
|  | description: predominant cell type | granulocytes, lymphocytes, macrophages, plasma cells, mixture (no predominant cell type) |
| 8 | necrotizing bronchitis, occurrence | 0 = not present, 1 = present in 1-3 foci, 3 = significantly present in >3 foci |
| 9 | necrotizing bronchitis, grade | for infiltrates /debris(1=few cells, 2 aggregates, 3 almost filling; 4 > filling lumen), grade max |
|  | description: predominant cell type | granulocytes, lymphocytes, macrophages, plasma cells, mixture (no predominant cell type) |
| 10 | Mucus increased | main = 1; periphery = 2 |
| 11 | infiltrates perivascular, occurrence | 0 = not present, 1 = present in 1-3 foci, 3 = significantly present in >3 foci |
| 12 | infiltrates perivascular, grade | 0 = no infiltrates, 1 = 1 cell layer, 2 = 2-3 layers, 3 = 4-5 layers, 4 = ≥6 layers, maximum score given |
|  | description: predominant cell type | granulocytes, lymphocytes, macrophages, plasma cells, mixture (no predominant cell type) |
| 13 | vascular rolling of immune cell, occurrence | 0 = not present, 1 = present in 1-3 foci, 3 = significantly present in >3 foci |
| 14 | endotheliitis, occurrence | 0 = not present, 1 = present in 1-3 foci, 3 = significantly present in >3 foci |
| 15 | vasculitis, i.e. mural damage, transmigrating immune cells | 0 = not present, 1 = present in 1-3 foci, 3 = significantly present in >3 foci |
| 16 | thrombi, occurrence | 0 = not present, 1 = present |
| 17 | necrosis, alveolar epithelium, area affected | 0 = no change, 1 = focal to oligofocal <5%, 2 = multifocal (6-40%), 3 = coalescing (41-80%), 4 = diffuse >80% |
| 18 | diffuse alveolar damage, with hyaline membrane, debris, fibrin, area affected | 0 = no change, 1 = focal to oligofocal <5%, 2 = multifocal (6-40%), 3 = coalescing (41-80%), 4 = diffuse >80% |
| 19 | hypertrophy/hyperplasia, bronchi, occurrence | 0 = not present, 1 = present in 1-3 foci, 3 = significantly present in >3 foci |
| 20 | hyperplasia /hypertrophy type II pneumocytes, area affected | 0 = no change, 1 = focal to oligofocal <5%, 2 = multifocal (6-40%), 3 = coalescing (41-80%), 4 = diffuse >80% |
| 21 | atypical cells or syncytia, occurrence | 0 = not present, 1 = present |
| sum scores | bronchial necrosis | sum of no. 8 to 9 |
|  | necrotizing bronchitis | sum of no. 6 to 9 |
|  | regeneration | sum of no. 19 to 20 |
|  | total lung score | sum of no. 1 to 21 |

\*scores can be translated to minimal (score 1), mild (score 2), moderate (score 3), severe (score 4)

